# H3K27me3 spreading organizes canonical PRC1 chromatin architecture to regulate developmental programs

**DOI:** 10.1101/2023.11.28.567931

**Authors:** Brian Krug, Bo Hu, Haifen Chen, Adam Ptack, Xiao Chen, Kristjan H. Gretarsson, Shriya Deshmukh, Nisha Kabir, Augusto Faria Andrade, Elias Jabbour, Ashot S. Harutyunyan, John J. Y. Lee, Maud Hulswit, Damien Faury, Caterina Russo, Xinjing Xu, Michael J. Johnston, Audrey Baguette, Nathan A. Dahl, Alexander G. Weil, Benjamin Ellezam, Rola Dali, Mathieu Blanchette, Khadija Wilson, Benjamin A. Garcia, Rajesh Kumar Soni, Marco Gallo, Michael D. Taylor, Claudia L. Kleinman, Jacek Majewski, Nada Jabado, Chao Lu

## Abstract

Polycomb Repressive Complex 2 (PRC2)-mediated histone H3K27 tri-methylation (H3K27me3) recruits canonical PRC1 (cPRC1) to maintain heterochromatin. In early development, polycomb-regulated genes are connected through long-range 3D interactions which resolve upon differentiation. Here, we report that polycomb looping is controlled by H3K27me3 spreading and regulates target gene silencing and cell fate specification. Using glioma-derived H3 Lys-27-Met (H3K27M) mutations as tools to restrict H3K27me3 deposition, we show that H3K27me3 confinement concentrates the chromatin pool of cPRC1, resulting in heightened 3D interactions mirroring chromatin architecture of pluripotency, and stringent gene repression that maintains cells in progenitor states to facilitate tumor development. Conversely, H3K27me3 spread in pluripotent stem cells, following neural differentiation or loss of the H3K36 methyltransferase NSD1, dilutes cPRC1 concentration and dissolves polycomb loops. These results identify the regulatory principles and disease implications of polycomb looping and nominate histone modification-guided distribution of reader complexes as an important mechanism for nuclear compartment organization.

**Highlights:**

□ The confinement of H3K27me3 at PRC2 nucleation sites without its spreading correlates with increased 3D chromatin interactions.
□ The H3K27M oncohistone concentrates canonical PRC1 that anchors chromatin loop interactions in gliomas, silencing developmental programs.
□ Stem and progenitor cells require factors promoting H3K27me3 confinement, including H3K36me2, to maintain cPRC1 loop architecture.
□ The cPRC1-H3K27me3 interaction is a targetable driver of aberrant self-renewal in tumor cells.

## Introduction

Nuclear activities are compartmentalized into various membraneless condensates, which contribute to partitioning chromatin events at specific loci in a precise spatiotemporal manner^1^. Whereas the biophysical mechanisms underlying formation and maintenance of nuclear condensates are increasingly appreciated^2,3^, there is limited understanding of how these compartments, often involving long-range 3D chromosomal interactions, are dynamically regulated during cell state transitions, and implicated in human diseases.

Polycomb Repressive Complexes 1 and 2 (PRC1, PRC2) are essential and conserved chromatin modifiers that regulate developmental cell fate transitions through cooperatively forming facultative heterochromatin^4,5^. PRC2 acts as the writer that catalyzes histone H3 lysine 27 mono-, di- and tri-methylation (H3K27me1/2/3) post-translational modifications (PTMs). PRC1 complexes contain a RING1A/B core subunit which catalyzes histone H2AK119 mono-ubiquitination (H2AK119ub) and can be further distinguished into canonical (cPRC1) or variant (vPRC1) complexes based on accessory subunits^6,7^. Polycomb bodies, identified by imaging studies over two decades ago, refer to nuclear foci of concentrated PRC1/2 complexes in association with H3K27me3 heterochromatin^8,9^. Through 3D genomics technologies, higher-order interactions between distal polycomb target regions have been described in several contexts of early development, including progenitor cells of embryonic and somatic tissues^5,10–19^. Intriguingly, polycomb bodies and polycomb-mediated chromatin loops are lost as differentiating cells exit stem/progenitor states^20,21^. The regulatory mechanisms and functional significance of this developmental stage-specific formation and dissolution of spatial 3D polycomb contacts remain unclear.

In mammalian cells, PRC2 nucleates at CpG Islands (CGIs) devoid of 5-methylcytosine (5mC) and can induce propagation of H3K27me1/2/3 over adjacent chromatin^22,23^. The spreading of H3K27me3 is controlled by regulatory stimuli and sensing of the local chromatin environment, including antagonistic histone PTMs^24^. For example, H3K36me2/3 and 5mC contribute to impairing PRC2 spread and demarcating boundaries between active euchromatin and H3K27me3 heterochromatin domains^23,25^. As such, the H3K36 dimethylase NSD1 is critical in several cell types for establishing intergenic H3K36me2 that serves to limit H3K27me3 spread^26,27^. As a result of tightly regulated PRC2 activity, genome-wide patterns of H3K27me3 are highly cell type-specific. Stem cells in primed pluripotency (PSCs) carry focal H3K27me3 confinement at a subset of promoters bearing a H3K4me3 co-enriched bivalent signature^28^, whereas in differentiated cells H3K27me3 can cover Mb-sized genomic regions. While it has been hypothesized that H3K27me3 spreading from nucleation sites may silence some adjacent promoters, repetitive elements and enhancers, the physiological importance of restrained versus broad H3K27me3 domains during lineage specification is incompletely understood.

Multiple cancers and genetic disorders are caused by mutations altering the quantity and genomic distribution of H3K27me3^29^. Diffuse midline pediatric high-grade gliomas (pHGGs) and posterior-fossa group A ependymomas (PFA-EPN) are deadly pediatric brain tumors defined by histone H3 Lys-27-Met substitutions (H3K27M), or by ectopic expression of EZH Inhibitory Protein (EZHIP) that shares structural resemblance to H3K27M^30,31^. This H3K27M oncohistone and its mimic EZHIP converge in mechanism to dramatically restrict H3K27me3 spreading beyond PRC2 nucleation sites, via allosteric impairment of EZH2/1 catalytic activity and PRC2 spatial redistribution. These effects cause global loss of H3K27me3 abundance in cell types otherwise capable of PRC2 spreading^32,33^. H3K27M/EZHIP impair glial cell differentiation and potentiate oncogenic transformation of progenitor cells in restricted developmental windows during early childhood^34–36^. Tumors harboring these alterations critically depend on residual H3K27me3^37,38^, distinguishing H3K27M/EZHIP cancers from those displaying complete loss of PRC2 function.

In this study, we use disease models with enforced H3K27me3 confinement (H3K27M/EZHIP-expressing tumors) or H3K27me3 expansion (NSD1 loss) as tools to identify a causal link between H3K27me3 patterns and 3D organization of polycomb target sites. We also investigate the mechanisms and impact of dysregulated polycomb condensation on silencing of polycomb target genes, cell differentiation and disease pathogenesis. Our results delineate how altered H3K27me3 spreading impacts chromatin architecture, gene expression and developmental transitions, and inform how these effects can promote disease states.

## Results

### Confined H3K27me3 spreading associates with enhanced 3D chromatin looping

To assess the relationship between 3D chromatin architecture and H3K27me3 confinement, we generated chromatin conformation capture (Hi-C) profiles from primary H3K27M pHGG (n=4) and matched pairs of patient-derived pHGG cell lines either bearing the H3.3K27M mutations, or in which we employed CRISPR-Cas9 editing to knock out the mutant H3.3K27M allele (KO, n=3), as previously described^39–41^. As controls, we also generated Hi-C datasets on other pediatric brain tumors (HGG wild-type for H3 mutations n=2, medulloblastoma n=9, ependymomas n=13, low-grade gliomas n=3) and normal midline brain tissue (n=3) (**Figure S1A-B**). Hi-C matrices at multiple scales, including compartment scores, topologically associating domain (TAD) boundary scores and matrix similarities, did not segregate H3K27M mutant from histone WT brain tumors (**Figure S1A-B**). When comparing H3K27M-mutants with matched KO control lines, the number of CTCF ChIP-seq peaks and contact frequencies among these sites were similar (**Figure S1C-D**), as were scores evaluating chromatin compartmentalization, insulation, and TAD architecture (**Figure S1E**).

Interestingly, we observed that H3K27M pHGG lines displayed distinctive long-range interactions between H3K27me3-marked regions (**Figure 1A**). Notably, frequencies of these 3D interactions anchored at H3K27me3 sites were significantly weakened upon H3K27M KO (**Figure 1A**). These structures can span up to tens of megabases, cross TAD boundaries, and anchor multiple sites at one location, in contrast to typical loop extrusion-associated structures that are generally on a sub-megabase scale^12,42,43^. These results indicate that despite broad conservation of overall 3D genome architecture, the confinement of H3K27me3 in H3K27M mutant tumors promotes the maintenance of 3D chromatin interactions with stereotypical features of polycomb bodies.

**Figure 1.**
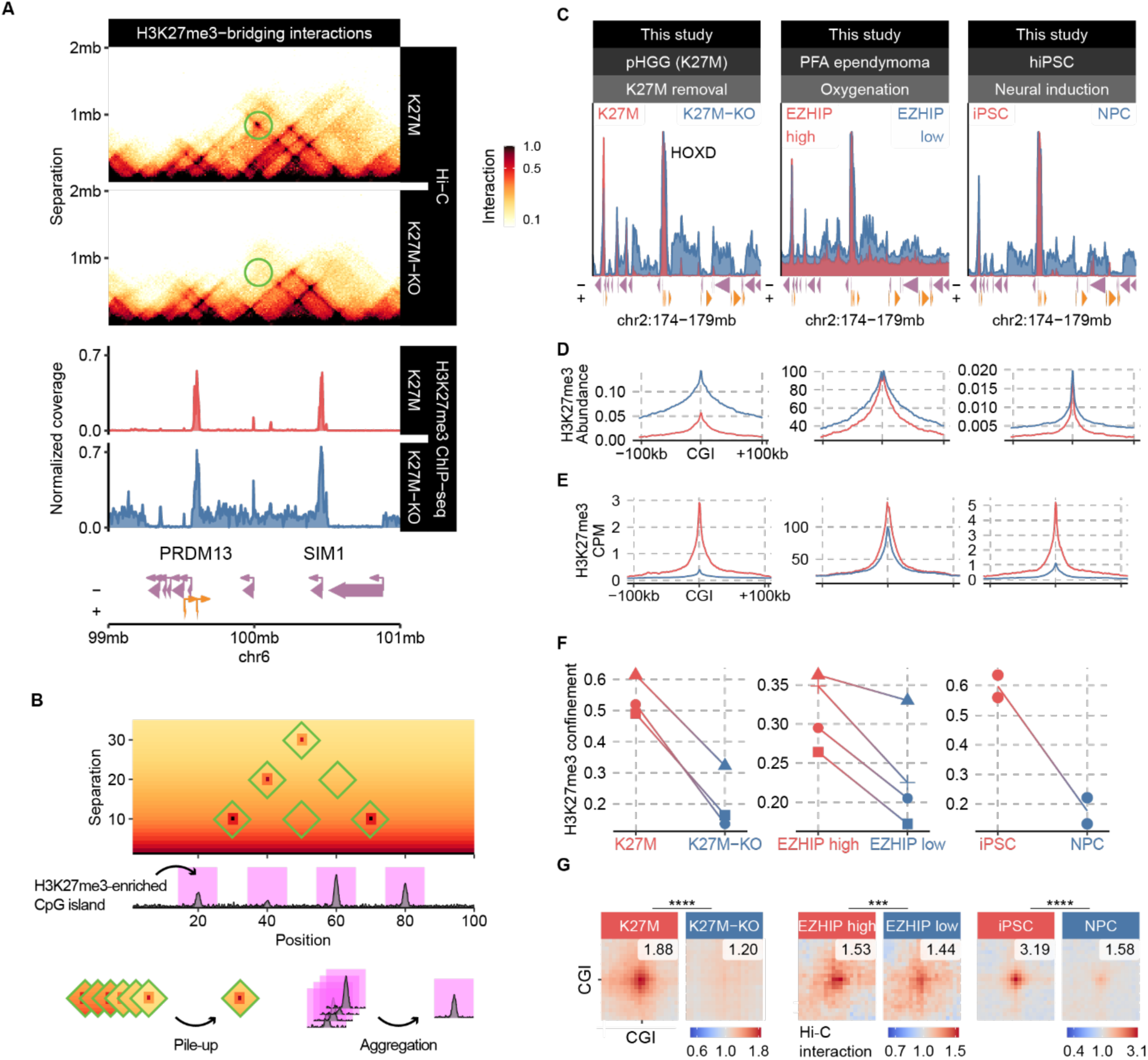
Restricted H3K27me3 occurs in brain tumor and developmental contexts and associates with greater frequencies of distal interactions between polycomb-marked CpG islands. A. Chromatin conformation capture (Hi-C) matrices showing a representative loop interaction (green circle) in a pHGG cell line BT245 (H3K27M versus H3K27M-KO) between the H3K27me3-enriched promoters of genes *PRDM13* and *SIM1* (top). ChIP-seq tracks of H3K27me3 showcase H3K27M-induced confinement of H3K27me3 at these genes’ promoters, whereas the spread of this mark covers a broad domain upon removal of the mutation in KO cells (bottom). B. Summary of quantitative approaches to measure aggregate enrichment of H3K27me3 at CGIs, and pile-up of pairwise contacts between H3K27me3-enriched CGIs in Hi-C data. C. ChIP-seq tracks of H3K27me3, normalized by Rx spike-in or abundance measured by mass spectrometry. A representative polycomb target locus (*HOXD* cluster) in isogenic matched comparisons of confined H3K27me3 due to H3K27M, EZHIP, or primed pluripotency is showcased. Loss of glioma drivers or iPSC-to-NPC differentiation is accompanied by the expansion of H3K27me3 domains in cell-type specific patterns. D. Metaplots of H3K27me3 aggregate ChIP-seq signals around H3K27me3-enriched CpG islands, normalized by total H3K27me3 abundance measured by ChIP-Rx spike-in. E. Metaplots as in D, normalized by read depth (counts per million; CPM). F. Measure of H3K27me3 ChIP-seq signal confinement (fragment cluster score at 1kb separation, computed using the tool “ssp”, see methods), comparing confined (H3K27M, EZHIP, primed pluripotency) versus diffuse profiles. Individual data points correspond to a replicate, with connected points indicating replicates from the same batch; connections not linking points indicate that multiple replicates were sequenced in a batch, and so the links are drawn between the average value per condition. G. Pile-up of Hi-C interactions among H3K27me3-enriched CpG islands, as defined in D, portraying average pairwise contact strength between such regions (in units of enrichment, i.e., observed / expected). Punctate enrichment signal in the center indicates elevated long-range interaction anchored at H3K27me3-enriched CGIs in cells with confined H3K27me3. H3K27me3-enriched is defined as the union set of top 1000 CpG islands with the most H3K27me3 alignments in either condition.

We next sought to address the nature of these 3D structures and their genome-wide relationship with H3K27me3 patterns. To this effect, we integrated H3K27me3 Chromatin Immunoprecipitation/Immunocleavage (ChIP/ChIC-seq) and Hi-C data from an additional brain tumor model with confined H3K27me3 deposition; EZHIP-expressing PFA-EPN, and a human induced pluripotent stem cell line (hiPSC, NCRM1) we manipulated to exit pluripotency by promoting its differentiation into neural progenitor cells (NPCs), a transition resulting in H3K27me3 spread (**Figure 1B-C**). Expression of H3K27M-like EZHIP in PFA-EPN derived cell lines is maintained in hypoxic culture conditions and largely decreased in normoxia, promoting spread of the H3K27me3 mark^37^. We thus generated datasets from PFA-EPN cell lines in hypoxia and normoxia (n=4) and from pluripotent hiPSCs and their derived NPC progeny (**Figure 1C**).

When applying ChIP-Rx (exogenous reference spike-in) normalization to H3K27me3 aggregate enrichment plots, H3K27M and EZHIP appear to diminish quantities of H3K27me3 at nucleating CGIs compared to isogenic controls where these genetic alterations are absent, similar to hiPSCs prior to their transitioning to NPCs (**Figure 1D**). However, when normalizing the same datasets by read depth (CPM, counts per million), the relative enrichment of H3K27me3 at these CGIs is greatly increased (**Figure 1E**), reflecting the confinement of residual H3K27me3 deposited at these loci. In order to derive quantitative measures of H3K27me3 confinement independent of its global quantity, we adapted a ChIP-seq quality control measure called Fragment Cluster Score (FCS) intended to assess the concentration of reads mapping to forward and reverse strands with different genomic separation^44^ (see Methods for additional details). This metric accurately established the dichotomy of confined versus diffuse H3K27me3 patterns based on simulating H3K27me3 profiles of varying breadth (**Figure S1F-H**). Importantly, the loss of H3K27M/EZHIP expression and hiPSC-to-NPC differentiation markedly reduced H3K27me3 confinement scores (**Figure 1F**). When applied to published ChIP-seq data from mouse embryonic brain^45^, this metric showed decreased H3K27me3 confinement with developmental age (**Figure S1I**), indicating that H3K27me3 domain expansion is observed during cellular differentiation *in vivo*.

Based on Hi-C data, we quantified the aggregate pair-wise interaction frequencies between H3K27me3-marked CGIs (**Figure 1B**). This approach allowed us to capture the enrichment of polycomb looping relative to background contact frequencies, which were significantly elevated by H3K27M/EZHIP expression and in primed pluripotency state (**Figure 1G**).

### Confined H3K27me3 spatially concentrates cPRC1 located on chromatin

To investigate the mechanisms behind H3K27M-induced polycomb loop formation, we charted the effects of H3K27M on PRC1 and PRC2 chromatin distributions and catalytic activity, using our isogenic pHGG lines. We used mass spectrometry (MS) to unbiasedly survey the presence and relative abundances of PRC1/2 subunits in nuclear soluble (nucleoplasm) and chromatin-bound protein fractions (**Figure 2A**). Both variant and canonical PRC1 members could be identified in chromatin-bound proteomes (**Figure S2H**). The pool of chromatin-associated cPRC1 contained varying abundances of CBX8, CBX2 and CBX4 subunits, and PHC1-3 subunits (**Figure 2B**). Despite ∼5-fold changes in H3K27me3 abundance, relatively minor differences in RNA expression (**Figure S2A**) and chromatin-bound quantities (**Figure 2B**) of CBX2, CBX4, and CBX8 were found between H3K27M and KO cells, using intensity-Based Absolute Quantification (iBAQ) measurements. Therefore, H3K27M mutations do not appreciably affect the composition and chromatin abundance of cPRC1 subunits.

**Figure 2.**
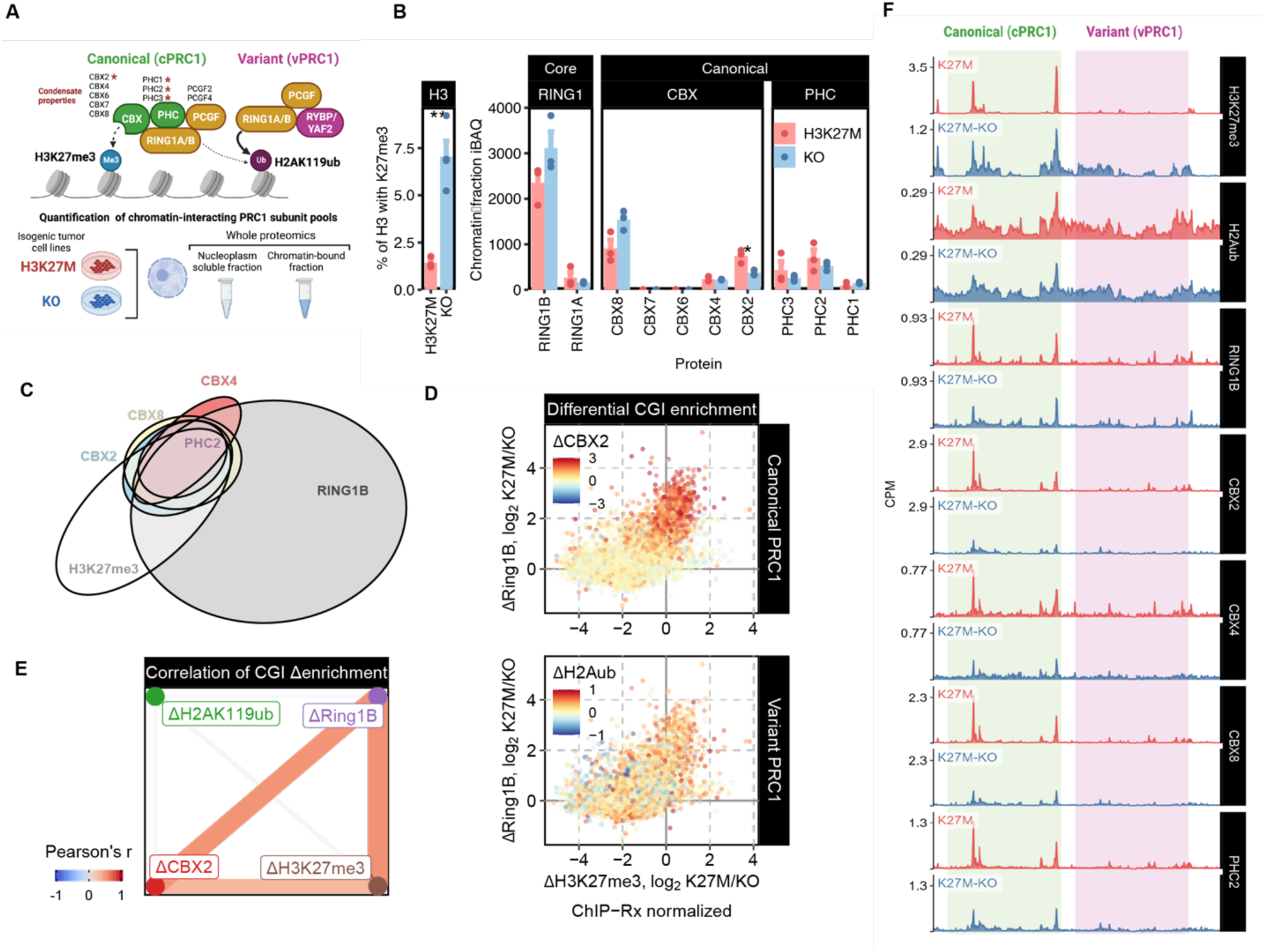
H3K27me3 confinement concentrates the chromatin pool of canonical PRC1. A. Schematic summary of PRC1 subunit composition defining core (yellow), canonical (green) and variant (pink) subcomplexes. Those known to engage self-associating properties are labeled with a star. B. Intensity-Based Absolute Quantification (iBAQ) values showing PRC1 subunit abundance in chromatin-fractionation lysates from DIPGXIII H3K27M and KO lines. H3K27me3 abundance (mass spectrometry) is included for comparison (left). C. Euler diagram of called peaks for H3K27me3, cPRC1 subunits (CBX2/4/8, PHC2) and core subunit RING1B in H3K27M pHGG cell line BT245. The strong overlap between CBX2/4/8 and PHC2 identify a common target group of cPRC1 sites, representing a minority of all PRC1 sites indicated by RING1B peaks. D. Density plots showing differential CGI enrichment of H3K27me3 (x-axis), RING1B (y-axis), and CBX2 or H2AK119ub (color code) between H3K27M and H3K27M-KO BT245 cells. Each dot represents a CGI and the differential enrichment is plotted as log2 ratio of K27M/KO. Retainment of H3K27me3 enrichment at CGIs associates with several fold greater enrichment for RING1B and CBX2 ChIP-seq signals, indicating the correlation between H3K27me3 confinement and enhanced cPRC1 recruitment (top). In contrast, there is a lack of correlation between H3K27me3 confinement and H2AK119ub enrichment (bottom). E. Correlation network of differential H3K27me3, RINGB1, CBX2 and H2AK119ub enrichment at CGIs of BT245 cells, demonstrating the weak correlation between H2AK119ub changes and changes of H3K27me3, RINGB1, CBX2. Edgewidths reflect the absolute value Pearson correlation coefficients. F. ChIP-seq tracks of a representative locus including cPRC1 sites (green) and adjacent vPRC1 site (pink). Track normalization by MS-iBAQ values provides quantitative measures of chromatin occupancy for each cPRC1 subunit, showing their concordance and heightened concentration due to the H3K27M mutation.

We next profiled the chromatin distributions of RING1B, CBX2/4/8 and PHC2 (most abundant PHC) cPRC1 subunits, and H2AK119ub (PRC1 catalytic product) by ChIP/ChIC-seq. Using peak calling, we identified a group of H3K27me3-enriched CGIs that significantly overlapped with RING1B binding sites (1686/3221, 52%), most of which (1148/1686, 68%) also coincided with CBX2/4/8 and PHC2 peaks (**Figure 2C**). This indicated that cPRC1 complexes, regardless of the constituent CBX subunit, co-localize to a subset of H3K27me3 sites. The majority of RING1B peaks (∼87%) were found outside of H3K27me3 domains, which likely represent binding sites of vPRC1 (**Figure 2C**). We next quantified how H3K27me3 spread affects cPRC1 chromatin localization. To this end, we used MS iBAQ measurements to normalize ChIP/ChIC-seq profiles of corresponding cPRC1 subunits. In H3K27M-mutants, CGIs retaining punctate H3K27me3 carried several-fold higher enrichment of RING1B and CBX2/4/8 compared to KO lines (**Figure 2D-F, S2B-E**). However, restriction of cPRC1 to these CGIs did not appreciably alter the abundance or profiles of H2AK119ub (**Figure 2D-F, S2D, F**), corroborating observations that this PTM is largely deposited by vPRC1 complexes in somatic cell types^46^. Importantly, we observed that the expansion of H3K27me3 domains in KO cells correlated with broad distributions of CBX2/8 and PHC2 enrichment at Mb-scale (**Figure S2G**). Therefore, genome-wide binding of cPRC1 closely follows H3K27me3 enrichment, which in H3K27M mutant cells effectively concentrates CGI promoter-proximal pools of the cPRC1 complex.

### Concentrating cPRC1 drives pathological polycomb condensates in H3K27M gliomas

We next sought to determine if the pathological formation of polycomb 3D interactions in H3K27M gliomas, resembling primed pluripotency, is mediated by aberrant concentration of cPRC1. To this end, we incorporated additional ChIP/ChIC-seq datasets of histone PTMs (H3K27ac, H3K4me3, H3K36me2/3, H3K9me3), architectural proteins (CTCF, cohesin complex member SMC1) and nascent transcription from Precision Run-On sequencing (PRO-seq) in three H3K27M pHGG lines (BT245, DIPGXIII, HSJ019) and clustered promoters and CGIs based on their chromatin states. We considered genomic intervals (promoters & CGIs) as individual observations and applied dimension reduction (UMAP) to summarize diverse epigenetic signals followed by unsupervised clustering (HDBSCAN) to derive discrete states. We identified four main groups; Cluster 1: Active (H3K27ac, H3K4me3, no repressive PTMs); Cluster 2: cPRC1 (CBX2/4/8, PHC2, RING1B, H3K27me3); Cluster 3: PRC2-only (H3K27me3, SUZ12, no PRC1) and Cluster 4: Other (lacking distinctive enrichment) (**Figure 3A-B, S3A-B**). We distinguished cPRC1 (Cluster 2) sites from the PRC2-only (Cluster 3) to test 3D structural changes associated with each complex. Using intra-class Hi-C pile up frequency measures (self-association), only Cluster 2 with cPRC1-marked regions displayed strong Hi-C interactions that resolved upon H3K27M KO, whereas PRC2-only sites were not affected (**Figure 3C**). cPRC1 clusters were also differentiated from PRC2-only clusters by lower average enrichment of CTCF signals (**Figure S3C**), consistent with cPRC1 interactions being independent of CTCF in embryonic stem cells^12^.

**Figure 3.**
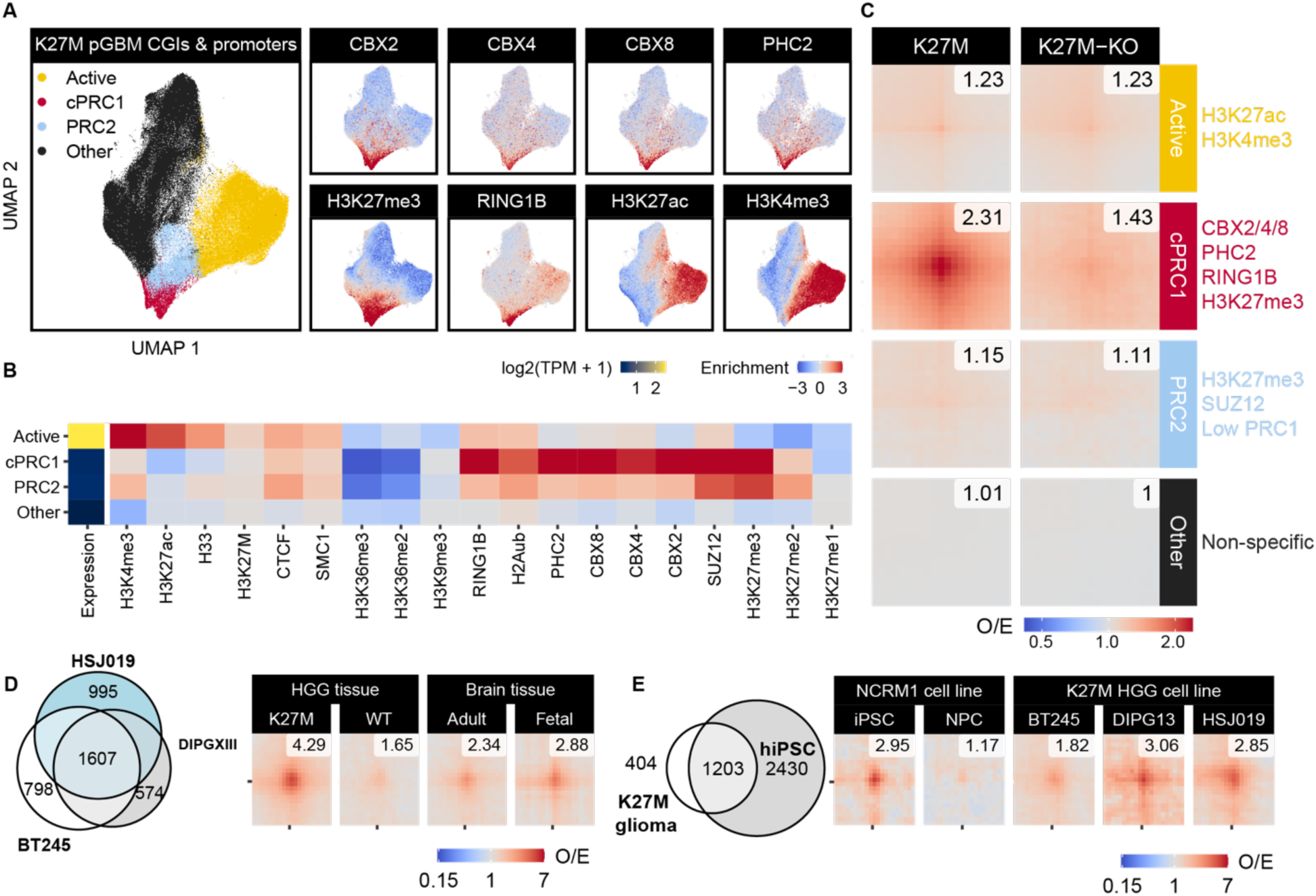
Concentrating cPRC1 drives pathological polycomb condensates in H3K27M gliomas. A. UMAP embedding and HDBSCAN clustering of chromatin state signals (ChIP/ChIC-seq enrichment) at CpG island & promoters (as listed in panel B) in the H3K27M line BT245 (all data from this study). Individual data points correspond to a genomic interval (promoter or CpG island), and the embedding is based on dimension reduction of features listed in panel B. Four different clusters/classes of regions are discovered: Active (enriched for H3K4me3 and H3K27ac), cPRC1 (with both PRC1 and PRC2 marks), PRC2-only (with SUZ12 and H3K27me3, lacking cPRC1), and Other (no specific enrichment). B. Average signals of transcription (RNA-seq, transcripts per million) and chromatin features for CGIs & promoters in different clusters, demonstrating the characteristic chromatin state of each cluster. C. Pile-up of Hi-C pairwise interactions were computed among genomic regions within the same cluster (i.e., intra-cluster loop contacts). cPRC1 cluster sites are the only cluster demonstrating heightened looping in H3K27M mutant but not KO lines of BT245. D. Euler diagram representing the overlap of cPRC1 cluster classification among 3 H3K27M cell lines (BT245, DIPGXIII, HSJ019) identifies 1607 consensus sites (left). Pile-up Hi-C pairwise interaction scores among consensus cPRC1 sites show substantially greater contact frequencies in H3K27M-mutant patient HGG tissues compared to histone WT HGGs and normal adult and fetal brain tissue, indicating that chromatin architecture differences observed in isogenic cell lines are conserved *in vivo*. E. The 1607 consensus cPRC1 sites substantially overlap with cPRC1 target sites in hiPSCs (NCRM1) derived from UMAP classification. Comparison of pile-up Hi-C pairwise interactions among the 1203 overlapping cPRC1 sites reveals H3K27M pHGG cell lines show strong cPRC1 looping, with interaction strength in a comparable range to iPSCs and substantially higher than iPSC-derived NPCs.

We observed substantial overlap of ∼1670 cPRC1-marked CGIs and promoters among the three cell lines derived from both thalamic and pontine H3K27M pHGGs that we designated as the consensus cPRC1 sites (**Figure 3D**). Functional analysis of these consensus cPRC1 sites revealed that they were enriched for genes involved in neural lineage differentiation pathways (**Figure S3D**) and that ∼75% sites were shared with those of the NCRM1 hiPSC line (**Figure 3E**). Furthermore, these shared cPRC1 sites showed comparable Hi-C aggregate 3D interaction scores between the iPSC line (2.95) and the three H3K27M pHGG lines (1.82-3.06). Last, similar to what we observed in H3K27M-KO cells, these scores decreased >50% (to 1.17) upon H3K27me3 spread and differentiation into NPCs (**Figure 3E**). These results suggest that, by confining H3K27me3 and preferentially concentrating cPRC1 to these confined sites, H3K27M maintains or recreates a higher order architecture that is transiently established by cPRC1 during early embryonic neural development. In agreement, in H3K27M patient tumor tissues, Hi-C interactions among consensus cPRC1 sites (4.29) were substantially higher than that in histone-WT pHGGs (1.65) and normal fetal or adult brain tissue (2.34-2.88) (**Figure 3D**). Therefore, H3K27M pHGG not only preserve or reproduce the cPRC1 landscape from early progenitors in their lineage of origin, they also actively maintain condensate-associated loop interactions at stronger intensity. This is achieved through limitation of H3K27me3 spread to PRC2 nucleation sites, thereby concentrating cPRC1 at a subset of these sites in nuclear space.

### H3K27me3 domain expansion weakens cPRC1 loop interactions in stem cells

The link between H3K27me3 confinement and cPRC1-mediated 3D interactions led us to investigate opposing scenarios where PRC2-mediated spreading of H3K27me3 is enhanced. Since H3K36me2 hinders PRC2 activity, the loss of this PTM can increase H3K27me3 spreading in stem cells and certain cancers^26,27,47^. As expected, *Nsd1* knockout (NSD1-KO) mouse embryonic stem cells (mESCs) showed a loss of H3K36me2 that correlated with global increase of H3K27me3 and its broad domain expansion throughout intergenic and genic regions^26^ (**Figure 4A**). Coinciding with H3K27me3 domain propagation, CBX2 binding was markedly diluted from focal peaks at H3K27me3-enriched CGIs upon *Nsd1* knockout (**Figure 4A-C**). We replicated our integrative UMAP classification of chromatin states at promoters and CGIs in this context and observed segregation of a class of cPRC1 targets carrying CBX2 enrichment (**Figure 4D, S4A**). Importantly, intra-class Hi-C aggregate interaction analysis revealed that *Nsd1* knockout specifically weakened 3D interactions among CBX2-containing cPRC1 sites, but not sites only enriched for PRC2 (**Figure 4D-E, S4A**). This data implicates a critical contribution of condensate scaffold CBX2 in bridging higher-order contacts in stem cells. The propensity for chromodomain-mediated CBX2 redistribution renders polycomb 3D looping sensitive to perturbations that abrogate H3K27me3 confinement, such as H3K36me2 loss.

**Figure 4.**
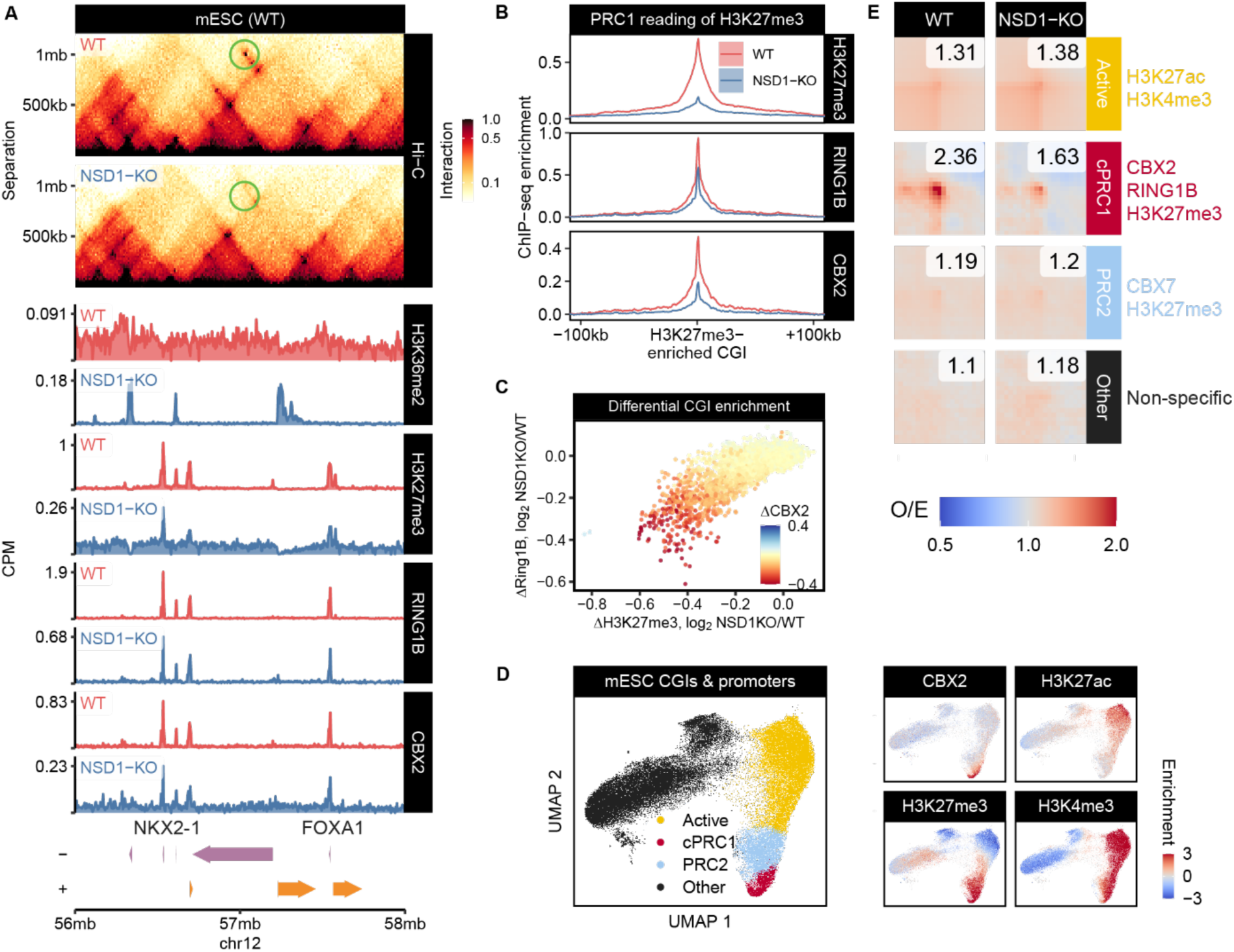
H3K27me3 spreading dilutes cPRC1 chromatin occupancy and weakens polycomb interactions in stem cells. A. Chromatin conformation capture (Hi-C) matrices showing a representative loop interaction (green circle) between cPRC1 sites bridging the promoters of *Nkx2-1* and *Foxa1* that is weakened upon KO of *Nsd1* in mESCs (top). ChIP-seq profiles of H3K36me2, H3K27me3, RING1B, and CBX2 (bottom) reveal that H3K27me3 spread accompanies H3K36me2 depletion in *Nsd1*-KO mESCs (blue), and cPRC1 binding becomes more diffuse compared to WT cells (red). B. Metaplot of H3K27me3 and PRC1 (RING1B, CBX2) aggregate ChIP-seq signal around H3K27me3-enriched CpG islands, in units of log2 enrichment over input, confirming *Nsd1*-KO reduces occupancy of cPRC1 at H3K27me3-enriched CGIs (union set of top 1000 most enriched in both conditions, as defined previously). C. Density plot showing differential CGI enrichment of H3K27me3 (x-axis), RING1B (y-axis), and CBX2 (color code) between WT and *Nsd1*-KO mESCs. Each dot represents a CGI and the differential enrichment is plotted as log2 ratio of Nsd1 KO/WT. Loss of CBX2 binding correlates with decreases in H3K27me3 and Ring1b at CGIs upon *Nsd1*-KO. D. UMAP embedding and HDBSCAN clustering of chromatin state signals at CpG islands and promoters in mESC (a combination of public and data from this study). Individual data points correspond to a genomic interval (promoter or CpG island), and the embedding is based on dimension reduction of all features. We derive four different clusters matching that of pHGG lines in Figure 3; Active, cPRC1, PRC2 (with SUZ12 and H3K27me3, lacking CBX2 enrichment), and Other. E. Pile-up of Hi-C pairwise interactions were computed among genomic regions within the same cluster (i.e., intra-cluster looping). On average, cPRC1 sites demonstrate the greatest differences in intra-class looping between WT and *Nsd1*-KO mESCs.

In addition to NSD1, several disease-associated genetic drivers can also alter H3K27me3 confinement^48^. Loss of linker histone H1 impairs H3K27me3 spread and facilitates lymphoma development^49^, and gain-of-function E1099K mutations in NSD2 drive shrinkage of H3K27me3 domains through expansion of H3K36me2 in acute lymphoblastic leukemias^50^ (among other types of cancer). Using published H3K27me3 ChIP-seq and Hi-C datasets, our analysis of confinement score and Hi-C pile-up frequencies found consistent correlations between H3K27me3 confinement and increased polycomb 3D interactions in models of H1-KO lymphomas and NSD2-E1099K-expressing leukemias (**Figure S4B-E**). These data reveal that the association between severely confined H3K27me3 and strengthened polycomb looping is found in multiple lineages in development and oncogenic transformation.

### cPRC1 loop architecture facilitates silencing of developmental gene expression programs

We next sought to determine the impact of changes in cPRC1 looping on transcription using transcriptome analysis by RNA-seq and PRO-seq (nascent RNA). In H3K27M cells, the 1607 Cluster 2 consensus cPRC1 sites (**Figure 3D**) could be further sub-clustered into two finer groups based on distinct chromatin states, designated sub-clusters A and B (**Figure 5A**). Hi-C pile-up interaction analysis revealed that sub-cluster A sites displayed higher and more focused aggregate contact frequencies in H3K27M mutants, whereas sub-cluster B sites showed weaker and more dispersed interactions (**Figure 5B**). The strong looping among sub-cluster A cPRC1 sites could also be observed in primary patient samples of H3K27M mutant pHGG and EZHIP-expressing PFA-EPN, compared to normal brain samples or other pediatric brain tumors (**Figure S5A**). In addition, sub-cluster A genes exhibited lower levels of promoter H3K4me3 and nascent transcription by PRO-seq (**Figure 5C**). Importantly, sub-cluster A genes displayed a significant, albeit moderate, upregulation of promoter and intragenic nascent RNA labeling in H3K27M KO lines, whereas PRO-seq signals at sub-cluster B genes remained unchanged (**Figure 5D**). Thus, strongly looped cPRC1 targets are more effectively silenced, and their repression is dependent on H3K27M-induced chromatin remodeling.

**Figure 5.**
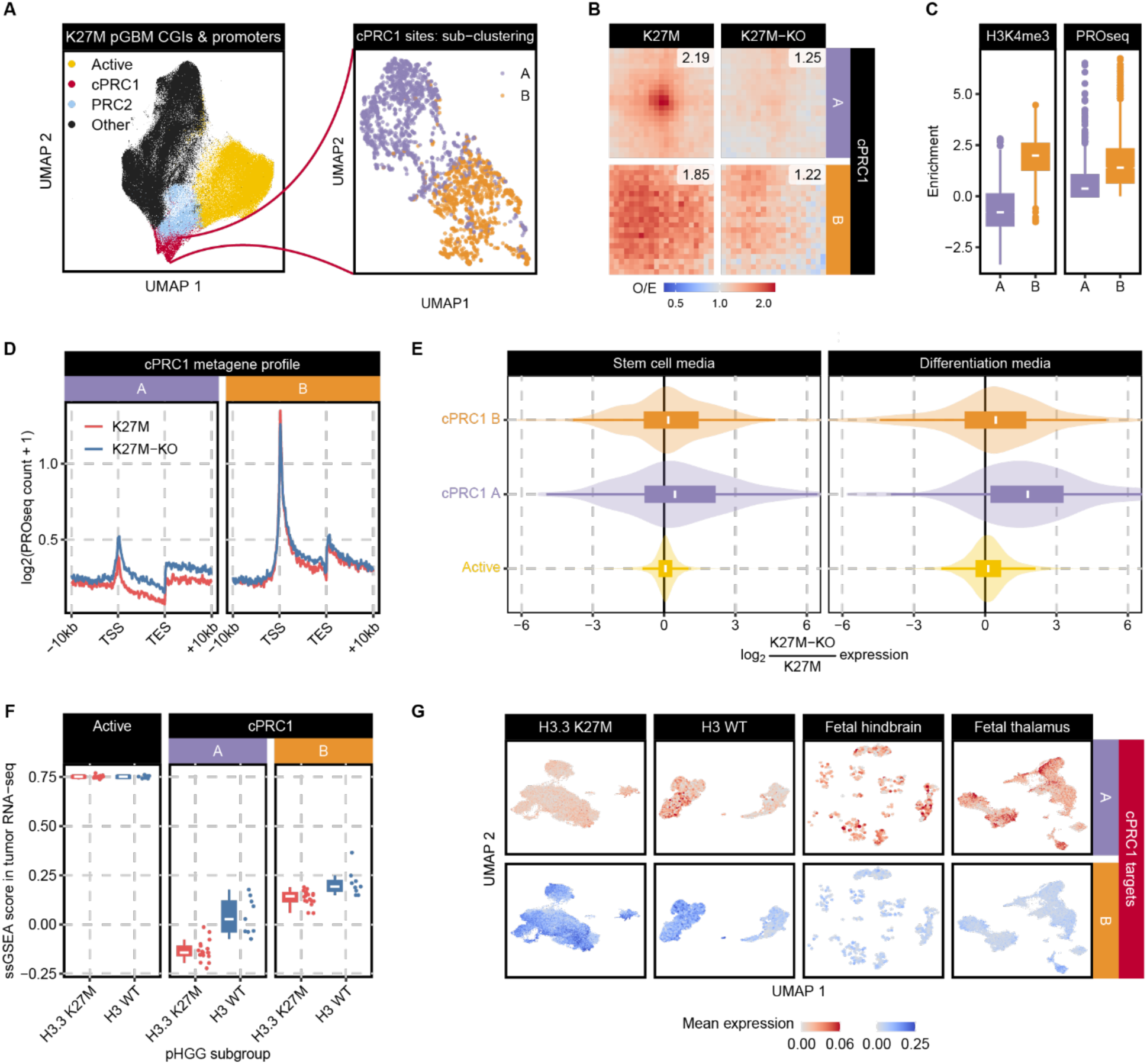
cPRC1 loop architecture facilitates silencing of developmental gene expression programs. A. UMAP-based subclustering for cPRC1 target sites, using the integration of all data (described in Figure 3), reveals two subclusters, denoted A and B. B. Pile-up of Hi-C pairwise interactions were computed among genomic regions within cPRC1 subclusters A and B. Subcluster A sites show more pronounced H3K27M-heightened loop interactions (focal intensity of signal enriched at the target site), whereas subcluster B sites show more diffuse interactions. C. Box plots of the enrichment scores for H3K4me3 and nascent transcription derived from PRO-seq experiments in H3K27M HGG cells. Subcluster A sites show significantly less enrichment of active chromatin state signals compared to subcluster B sites. D. Meta-gene plots of nascent transcription rates (PRO-seq signal) from subcluster A and B target genes in matched H3K27M and KO lines (BT245). H3K27M-KO enhances promoter and gene body nascent transcription of subcluster A targets, whereas subcluster B targets show substantially higher transcription, independent of the H3K27M mutation. E. Violin plots of genes’ differential expression values in RNA-seq data. cPRC1 subcluster A targets undergo greater upregulation than subcluster B upon H3K27M-KO. The effect size is moderate when cells are maintained in stem cell media (left), and becomes more pronounced when cells are cultured in differentiation media promoting glial maturation (right). F. Single sample gene set enrichment (ssGSEA) scores show that the representation of cPRC1 subcluster A genes is depleted in transcriptomes of H3K27M patient tumors compared to H3 WT pHGG tumors. In contrast, subcluster B and active genes are equivalently expressed in both groups, showing H3K27M tumors *in vivo* maintain strong repression of cPRC1-looping genes. G. UMAP projections of scRNA-seq data from H3K27M mutant or H3 WT HGGs, or normal brain tissues. Each single cell is colored based on their mean expression values for cPRC1 subcluster target genes. cPRC1 subcluster A gene expression is uniformly low across all H3K27M tumor cells, whereas these genes are variably expressed among cell populations of WT HGGs and fetal brain tissues. cPRC1 subcluster B genes are expressed comparably in both glioma groups.

Despite a modest increase in nascent transcription, sub-cluster A genes did not show substantial upregulation of steady-state mRNA abundance measured by bulk RNA-seq in H3K27M KO lines maintained in stem cell media (**Figure 5E, S5B**). We have previously shown that while H3K27M mutation impairs glial differentiation, this effect is only manifest when replacing stem cell media with those promoting glial maturation (differentiation media)^51^. Accordingly, expression of sub-cluster A targets were significantly upregulated in H3K27M KO compared to mutant lines after a 14-day course of differentiation (**Figure 5E, S5B**).

To extrapolate these findings into patient settings, we analyzed both bulk and single-cell RNA-seq datasets of pHGG primary tumors. Single set Gene Set Enrichment Analysis (ssGSEA) shows that sub-cluster A genes were substantially depleted in H3K27M patient tumor transcriptomes compared to WT pHGGs, whereas sub-cluster B genes were expressed at substantially higher levels irrespective of the mutation (**Figure 5F**). Furthermore, sub-cluster A gene representation was homogenously absent in H3K27M tumor single-cell RNA-seq (scRNA-seq) data across all tumor cell populations, whereas these same genes were variably expressed across neural progenitors, astrocytes, oligodendrocytes and neuronal populations of WT tumors and normal fetal brain (**Figure 5G, S5C**). Our data indicate that the 3D polycomb loops generated by restricting cPRC1 to H3K27me3 peaks promote stringent repression of gene expression at these loci, thereby providing a plausible mechanism for the known enhanced polycomb target repression by the H3K27M oncohistone^38,39,52,53^.

### cPRC1 looping contributes to the restricted developmental potential during H3K27M gliomagenesis

We next examined the functional significance of repressive cPRC1 looping using *in vivo* models of H3K27M tumorigenesis. We previously reported that H3K27M is required to maintain tumor-forming competence of pHGG cells engrafted in orthotopic sites of immunodeficient mice^39,54^. H3K27M mutant lines, capable of forming high-grade tumors in mice with high penetrance, experience loss or substantial delay of tumor formation when the mutation is removed. Despite the lack of tumor development in mice injected with H3K27M-KO pHGG lines, we found extensive seeding of human cells throughout the murine brain, using luciferase-labeled cell imaging and transcriptomics (**Figure 6A, S6A**). We recovered three matched pairs of engrafted H3K27M and KO cell lines (BT245, DIPGXIII, HSJ019) from mouse brains and performed scRNA-seq to map *in vivo* cell states to our previously published reference atlas of brain development^51^. H3K27M tumor cell populations displayed lower representations of mature glial cells and increased fractions of glial progenitor-resembling cells, including radial glial cells (RGCs) at the top of the neural development hierarchy (**Figure 6B-D, S6B**). In contrast, in H3K27M-KO cell populations, RGCs were lost or largely diminished in number. Instead, there were greater fractions of cells resembling mature astrocytes and oligodendrocytes, represented by the marker genes Glial Fibrillary Acidic Protein (*GFAP*) and Myelin Basic Protein (*MBP*), respectively (**Figure 6B-C**). The impaired differentiation states of H3K27M xenografts correlated with differential expression of sub-cluster A target genes, which were consistently repressed in H3K27M tumors, compared to their levels across all KO cell type populations (**Figure 6C, S6B**). Together with patient data, these results implicate a role for cPRC1 condensate and looping in limiting the expression of target genes involved in neural development, thereby contributing to restricted differentiation potential of H3K27M-mutant tumors.

**Figure 6.**
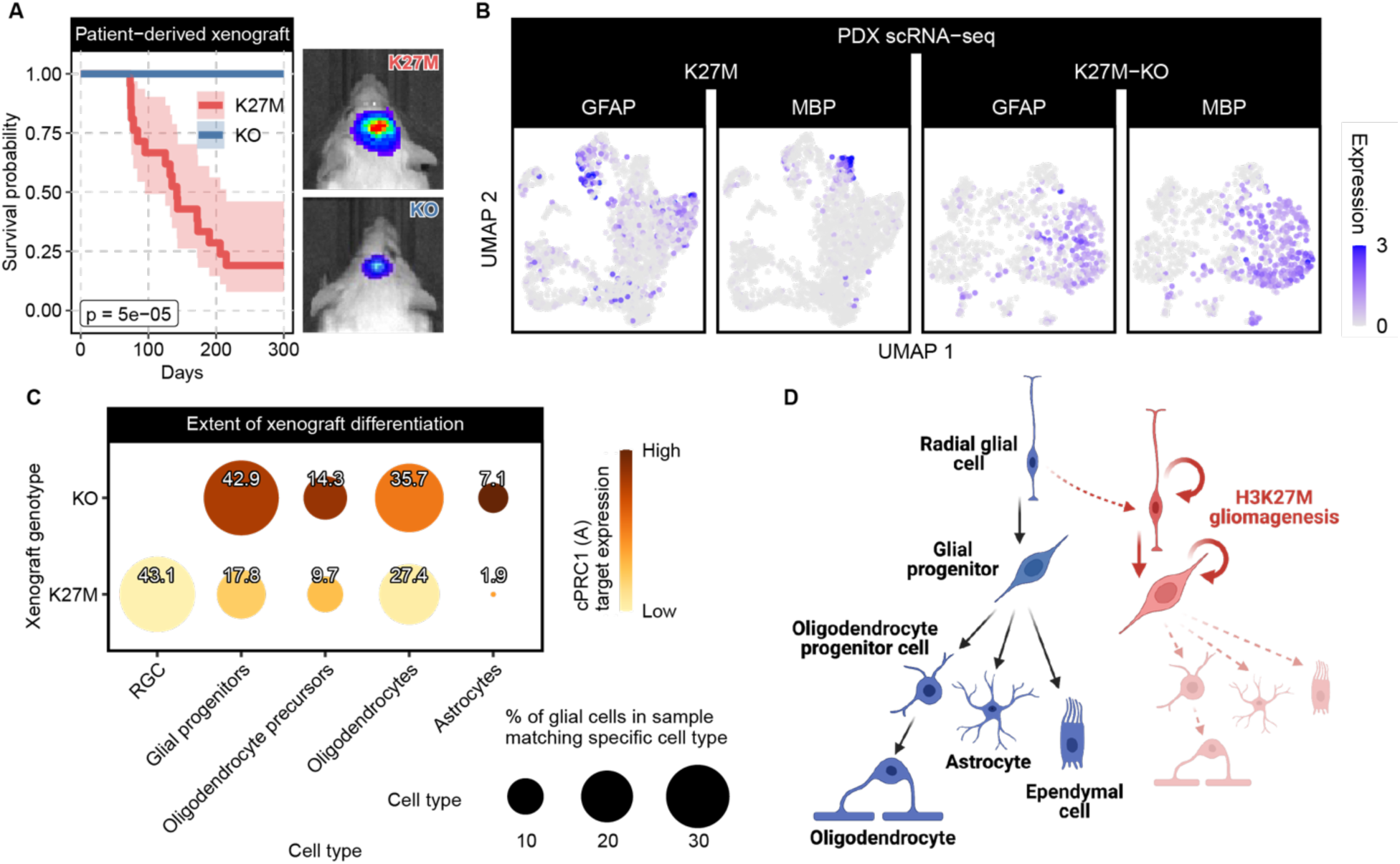
cPRC1 looping contributes to the restricted developmental potential during H3K27M gliomagenesis. A. Kaplan-Meier survival curves for H3K27M and KO pHGG cell lines (BT245) in orthotopic xenograft models identify a dependence on H3K27M to maintain tumorigenic potential *in vivo* (left) (with 95% confidence interval shaded). *In vivo* imaging of tumor cell luciferase signals shows representative tumor seeding at the caudate putamen injection site and greater signal intensity in lethal H3K27M tumors (right). B. UMAP projections of populations of tumor cells recovered and profiled by scRNA-seq. Each cell is colored based on expression levels of *GFAP* or *MBP*. H3K27M cells in xenograft tumors (left) show lower expression of the oligodendrocyte differentiation marker Myelin Basic Protein (MBP) and astrocyte marker Glial Fibrilary Acidic Protein (GFAP) than KO engrafted cells (right). C. Bubble plots representing fractions of tumor cell populations annotated by cell type classification (bubble size) based on ssGSEA mapping to reference cell types (see Methods). Each bubble is also colored based on the level of cPRC1 subcluster A target gene expression (bubble color). Expression of cPRC1 subcluster A genes is consistently depleted across the spectrum of classified cell types in H3K27M samples, accompanied by lower proportions of more differentiated cell types (astrocytes, oligodendrocytes). D. Schematic summary of developmental trajectories among neural cell types related to gliomagenesis, and the relative cell type transitions characterizing H3K27M (red) and KO (blue) xenograft cell populations. The schema is based on analysis of 3 cell line models (BT245, supplemental: DIPGXIII, HSJ019).

### Inhibiting CBX chromodomain-H3K27me3 interaction restores differentiation potential of H3K27M gliomas

cPRC1 residence time on chromatin relates to the affinity of its CBX subunits’ chromodomains for H3K27me3^55^. Therefore, we tested whether blocking the interaction between CBX chromodomain and H3K27me3 could affect cPRC1 condensation and looping, and in turn alter the differentiation potential of H3K27M pHGG cells. We applied UNC4976, a chemical probe termed CBX allosteric modulator (CBX-AM) that selectively obstructs CBX chromodomain reading of H3K27me3^56,57^, to three H3K27M pHGG lines. In BT245 cells, treatment of CBX-AM UNC4976 diluted chromatin enrichment of CBX2 and RING1B subunits from cPRC1 sites, whereas RING1B enrichment at active sites was unaffected (**Figure 7A**). This coincided with a decreased Hi-C aggregate interactions specifically at sub-cluster A sites, confirming that cPRC1 condensate and looping require a critical threshold of local complex concentration (**Figure 7B**). Also, RNA-seq showed stable upregulation of sub-cluster A genes by CBX-AM treatment in H3K27M pHGG lines following differentiation stimuli (**Figure 7C, S7A**). We next measured the acquisition of GFAP and SRY-Box Transcription Factor 10 (SOX10), markers of astrocyte and oligodendrocyte differentiation, respectively. Treatment of CBX-AM in H3K27M DIPGXIII or BT245 lines induced GFAP and SOX10 expression to levels comparable to those observed in KO lines (**Figure 7D, S7B-C**), signaling that by promoting cPRC1 loop dissolution we could endow greater competence for differentiation despite the presence of the H3K27M mutation.

**Figure 7.**
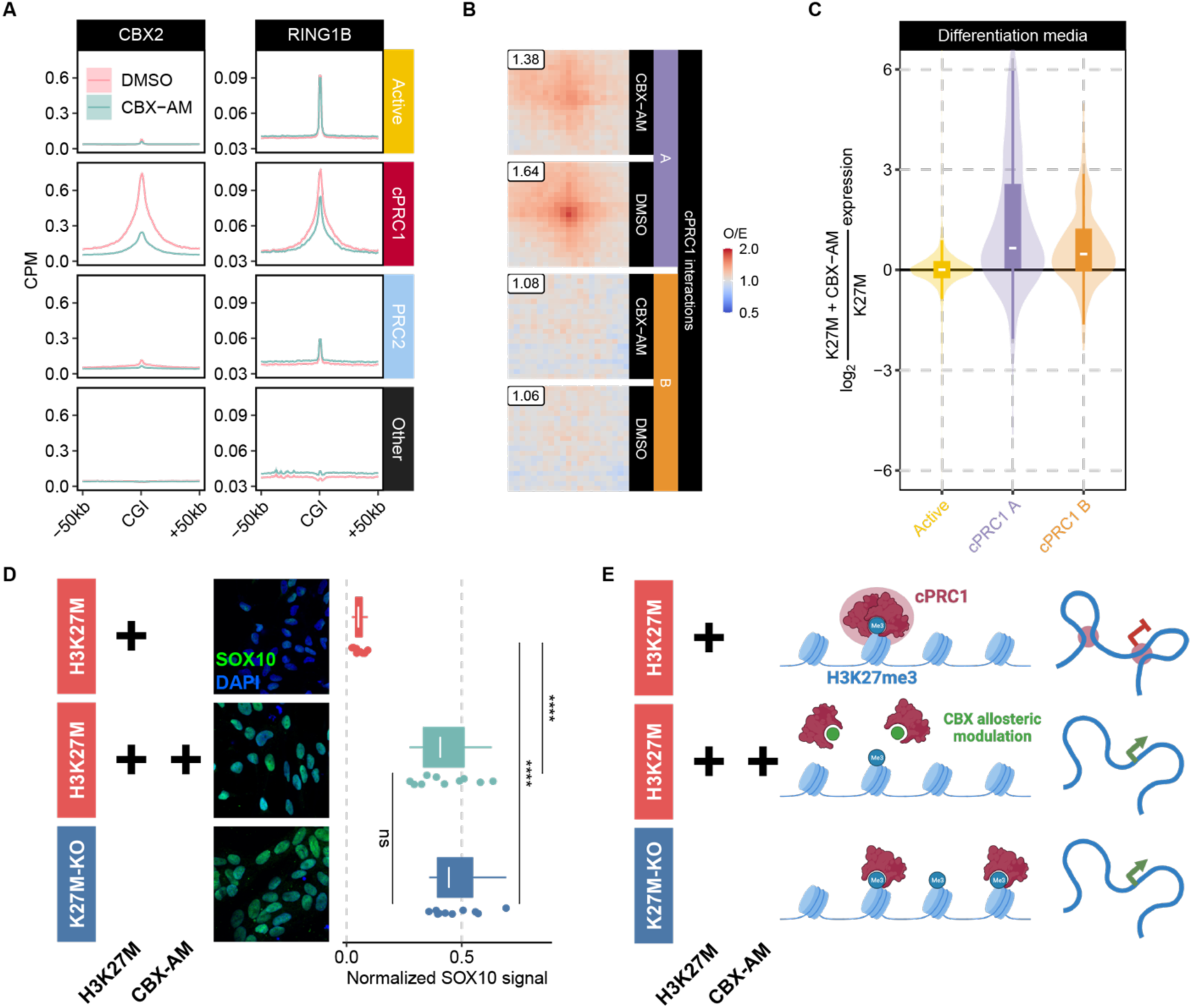
Blocking CBX chromodomain function relieves cPRC1 loop-associated repression and promotes tumor differentiation. A. Metaplots of CBX2 or RING1B aggregate ChIP-seq signals around CpG islands within each annotated cluster (see Figure 3), for BT245 H3K27M lines treated with DMSO control or CBX-AM compound. CBX-AM treatment attenuates the enrichment of RING1B and CBX2 at cPRC1 target sites. B. Pile-up of Hi-C pairwise interactions were computed among genomic regions within cPRC1 subclusters A and B, in BT245 H3K27M lines treated with DMSO or CBX-AM. CBX-AM treatment weakens frequencies of Hi-C interactions specifically at cPRC1 subcluster A sites. C. Violin plots of genes’ differential expression values in RNA-seq data. CBX-AM treatment results in upregulation of cPRC1 subcluster A genes, and to a less extent subcluster B genes, in H3K27M BT245 cells treated with differentiation media. Violin plots’ hinges correspond to the 25^th^ and 75^th^ percentiles, with whiskers extending to the most extreme value within 1.5 × interquartile range from the hinges, whereas the central band mark the median value. D. Immunofluorescence microscopy imaging of the oligodendrocyte progenitor cell differentiation marker SRY-box Transcription Factor 10 (SOX10). CBX-AM treatment elevates SOX10 expression in H3K27M BT245 cells to levels comparable to KO cells. E. Summary of CBX-AM treatment and H3K27M-KO’s effect on H3K27me3, CBX chromodomain localization and chromatin architecture of H3K27M pHGG cells. Manipulation of cPRC1 concentration through either H3K27M KO or CBX-AM dilution of chromodomains results in loss of repressive loop architecture and potentiation of differentiation.

We corroborated these findings made by chemical probes with a genetic approach using stable knockdown of CBX and PHC2 proteins. ShRNA-mediated depletion of both CBX2 and CBX4 consistently led to upregulation of sub-cluster A gene targets (**Figure S7D-E**). This indicates that multiple cPRC1 subunits could mediate looping-associated transcriptional repression in a cooperative and/or partially redundant manner. Taken together, H3K27M-mediated confinement of H3K27me3 drives the concentration of cPRC1 to a critical threshold required for loop formation and stable gene silencing. Dilution of the local chromatin pool of cPRC1, through either H3K27me3 spread, chromodomain obstruction or decreased CBX dosage, can alleviate target gene repression anchored in loop interactions, thereby restoring cells’ competence for lineage maturation (**Figure 7E**).

## Discussion

Polycomb bodies represent a major class of nuclear condensates mainly present in early development and specific cell types, which tether distal genomic regions in 3D. How these higher-order nuclear structures are formed is largely unknown while their functional relevance is being debated. Using H3K27M and EZHIP glioma models, we show that the substrate-reader relationship of H3K27me3 and chromodomains determines local concentrations of cPRC1 and controls the formation of these chromatin condensates. Furthermore, by modulating their genesis and dissolution, we identify the role of cPRC1 looping in enforcing repression of target polycomb genes during development and tumorigenesis.

Using isogenic tumor models including H3K27M-mutant, EZHIP-expressing and NSD1 KO lines, and iPSCs differentiated into NPC, we show that polycomb 3D architecture is dynamically regulated in early progenitors during cell state transition. Unlike studies *in vitro* using PRC1/2 overexpression, we show that the formation of these structures under physiological conditions occurs without changes in global levels of PRC1/2 components. Instead, two distinct patterns of H3K27me3 deposition guide the chromatin distribution and local concentration of cPRC1: In the context of stem cells and in tumor cells expressing the H3K27M mutation or EZHIP, H3K27me3 is focal, restricted to specific CGIs representing PRC2 nucleation sites. The relative enrichment of the mark at these loci is what promotes close to exclusive recruitment of its CBX readers and cPRC1. We propose that accumulation of CBX2 allows it to reach a critical threshold required for condensate formation, which would be mediated by the liquid-liquid phase separation properties of its intrinsically disordered region (IDR), as well as oligomerization of Sterile alpha motif (SAM) domains in Polyhomeotic-like proteins (PHC1/2/3)^58,59^. This in turn leads to the formation of polycomb bodies that can tether distant polycomb targets for strong and coordinated repression. As polycomb gene repression has been linked with diminished RNA polymerase II binding, pause-release and transcriptional burst frequencies^60^, future studies are needed to address whether, and how, condensation-mediated 3D clustering can limit the access of these transcriptional machineries to cPRC1 target genes.

Beyond providing a rationale for H3K27M and EZHIP effects in stalling differentiation in cancer, our findings more broadly suggest a mechanism for the resemblance between clinical phenotypes of Sotos Syndrome (germline loss-of-function heterozygous mutations in NSD1) and Weaver syndrome-related overgrowth disorders (germline mutations in EZH2, SUZ12 or EED subunits of PRC2)^61,62^. Spread of H3K27me3 from CGIs due to decreased H3K36me2 boundary in NSD1-mutant Sotos patients may lead to precocious dissolution of cPRC1 condensates in progenitor cells. Conversely, diminished H3K27me3 deposition following decreased PRC2 activity in Weaver patients may lead to lower recruitment of cPRC1 members below the critical levels needed for condensate and loop formation. In both diseases, untimely dissolution of cPRC1 condensates would result in early transcription of normally repressed polycomb targets and premature differentiation and lineage commitment, accounting for patients’ characteristic accelerated growth rates from early childhood and advanced morphological and molecular aging markers^63^. Notably, the Hox gene clusters are ancient polycomb targets that display altered chromatin in cancers including H3K27M pHGGs and in Sotos syndrome^64–66^. Variations in the potential of polycomb condensates to repress master regulatory Hox segments could contribute to either stalling H3K27M tumor differentiation or, in contrast, accelerating developmental clocks in overgrowth disorders. In addition, in a genetic screen for maternal-effect sterile (MES) mutants in *C. elegans*^67^, overlapping transcriptional effects are noted in mutants of MES-2, MES-3 and MES-6, which form the *C. elegans* PRC2 complex^68^, and in MES-4, which catalyzes broad domains of H3K36 methylation similar to NSD family enzymes^69,70^. Thus, the phenocopy between NSD and PRC2 mutations during development from worms to human suggest that the partitioning of H3K36 and H3K27 methylation domains may be an evolutionarily conserved mechanism to organize higher-order architecture and coordinate expression of polycomb target genes to regulate differentiation and lineage commitment.

In H3K27M pHGGs, chemical obstruction of chromodomain function can dissolve loop architecture, alleviate target gene silencing and promote differentiation. This approach may represent a potential therapeutic avenue in these deadly tumors and other cancers where polycomb looping likely plays an oncogenic role, including H1-mutant lymphomas and NSD2 gain-of-function leukemias. Targeting cancer cells’ pathologically heightened cPRC1 condensates could be an attractive therapeutic strategy that limits toxicity in differentiated somatic cell types where this nuclear compartment is less often observed.

In summary, our results identify the role of H3K27me3 spread in modulating polycomb target gene expression, which orchestrates developmental transitions through the maintenance or dissolution of cPRC1-mediated 3D interactions. Imbalances in the spread of this mark alter genome architecture and contribute to the disease mechanism of H3K27M gliomas and potentially other cancers and NSD1/PRC2-associated developmental syndromes. More broadly, this model represents a demonstration of how histone PTM codes, established by writer proteins that nucleate at regulatory elements and spread across genomic regions, subsequently segregate pools of reader complexes throughout the nuclear space. This spatial organization of writer-reader complex association can then partition active or repressive chromatin compartments at both local and distal scales through condensate formation, facilitating dynamic and coordinated gene regulation during cell state transitions.

## Acknowledgements

This work was supported by a Large-Scale Applied Research Project grant from Genome Quebec, Genome Canada, the Government of Canada, and Ministère de l’Économie, de la Science et de l’Innovation du Québec, with the support of the Ontario Research Fund through funding provided by the Government of Ontario to N.J., J.M. and C.L.K.; the Canadian Institutes for Health Research (CIHR; grants PJT-156086 to C.L.K., PJT-183939 to J.M., and MOP-286756 and FDN-154307 to N.J.); the National Sciences and Engineering Research Council (NSERC; grant RGPIN-2016-04911 to C.L.K.). J.M. and N.J. is supported by funding from the United States National Institutes of Health (NIH) Grant P01-CA196539; C. L. is supported by funding from NIH Grant R35GM138181 and the Pew-Stewart Scholars for Cancer Research Award; B. A. G. is supported by NIH grants CA196539 and AI118891, and the St. Jude Children’s Hospital Chromatin Consortium; B. H. is supported by the Canadian Institutes of Health Research (CIHR) Banting and Best Graduate Scholarship. C.L.K. is supported by a salary award from Fonds de recherche du Québec (FRQS). N.J. is a member of the Penny Cole Laboratory and holds a Canada Research Chair Tier 1 in Pediatric Oncology from CIHR. This work was performed within the context of the International Childhood Astrocytoma Integrated Genomic and Epigenomic (ICHANGE) consortium. Data analyses were enabled by compute and storage resources provided by Compute Canada and Calcul Québec. We are especially grateful for the generous philanthropic donation of the Charles Bruneau Foundation, the WeLoveYouConnie Foundation, and the Cedars/Sarah Cook funds.

## Author contributions

A. B. K. and B. H. generated data and contributed to study design, data interpretation and manuscript preparation. H. C., X. C., N. K., A. B., M. J. J., R. D. and M. B contributed to sequencing data analysis and interpretation. A. P., S. D., K. H. G., A. F. A., E. J., A. H., J. J. Y. L., M. H., D. F., C. R and X. X. contributed to data generation, analysis and interpretation. N. A. D., A. G. W. and B. E. assisted with the collection of patient samples, study design and data interpretation. R.K.S. led the interactome and global quantitative proteomic design, analysis, and interpretation. K. W. and B. A. G. led the histone proteomics data generation and analysis. M. G., M. D. T., C. L. K., J. M., N. J and C. L. contributed to study design, data interpretation, and manuscript preparation.

## Additional Information

Supplementary Information is available for this paper. Correspondence and requests for materials should be addressed to Nada Jabado.

---

**Supplemental Figure SF1.**
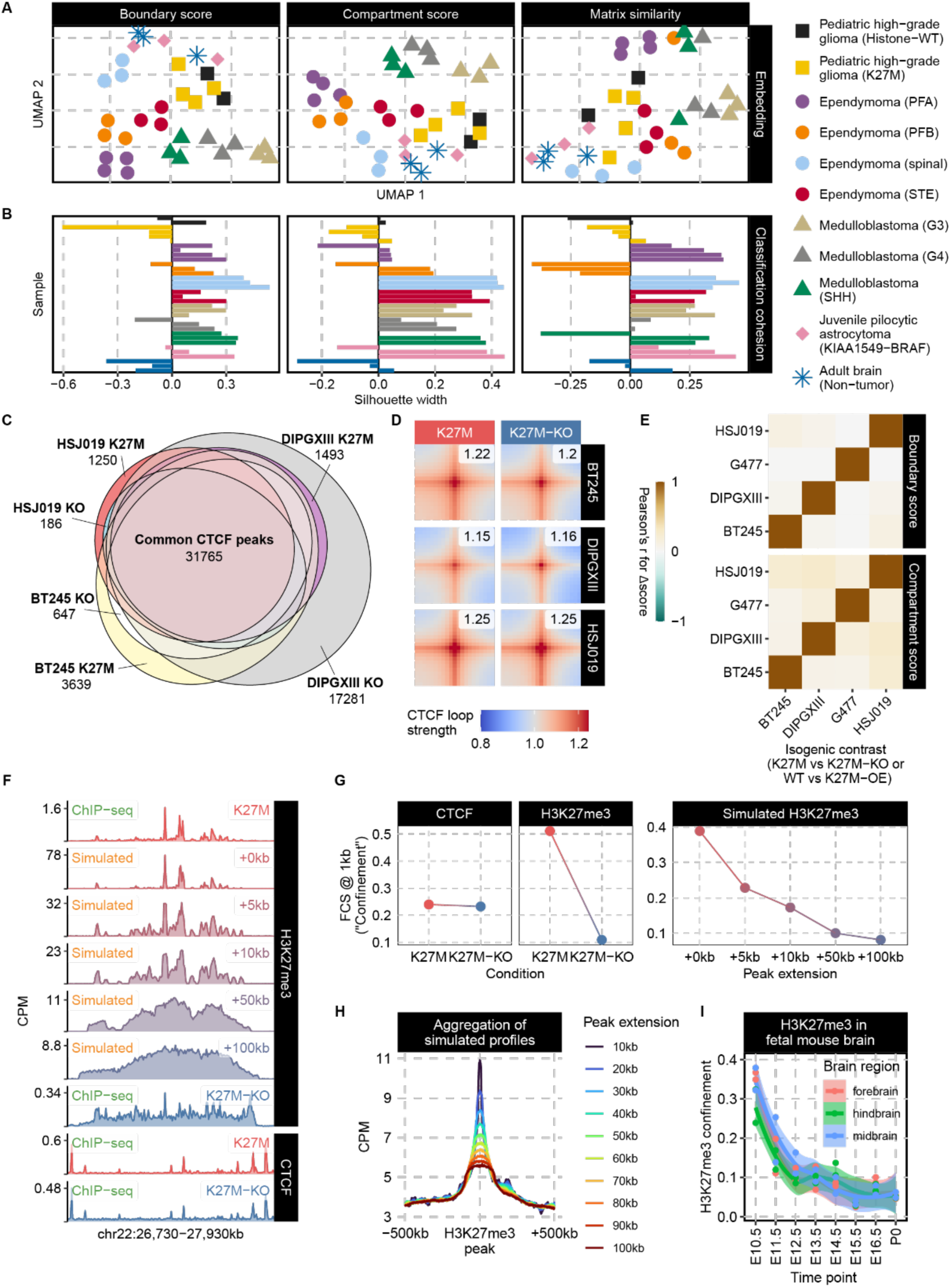
A. Hi-C data generated from brain tumors and normal brain tissues are analyzed at genome-wide scales. UMAP embedding based on genome-wide comparison across Hi-C contact matrices at three different scales: compartmentalization (first principal component / compartment score), topologically associating domain organization (RobusTAD boundary score), and matrix similarity (HiCRep coefficient). H3K27M pHGGs do not separate from H3 WT pHGGs by any of the three modalities examined. B. From Hi-C datasets in A, silhouette width based on inter-sample similarity in terms of three different modalities, with more positive values indicating that a sample is closer to other samples belonging to the same class whereas more negative samples indicating lack of cohesion (i.e., class label is not reflected by high inter-sample similarity for those belonging to the same class). H3K27M pHGGs emerge as the only tumor subtype demonstrating lack of distinct signatures across all three scales considered, generally showing negative silhouette scores (i.e., less similar to other H3K27M pHGGs than to tumors of another type). Therefore H3K27M does not impose a specific signature on large-scale genome organization. C. Euler diagram of CTCF peaks identified in isogenic H3K27M pHGG cell lines and their KO counterparts, demonstrating a substantial overlap. D. Pile-up of pairwise Hi-C interactions among the union CTCF peak set across all H3K27M and KO samples; only pairs of sites with convergent motif orientations were considered. This reveals a lack of global differences in CTCF interaction strength between isogenic H3K27M and H3K27M-KO pHGG cells. E. Correlation of compartment/insulation score differences (H3K27M versus KO/WT) between isogenic comparisons. The weak correlation coefficients demonstrate lack of consistent changes in compartment/domain structures upon the removal or overexpression of H3K27M. F. Representative tracks of experimental and simulated ChIP-seq datasets, demonstrating the distinction between confined versus diffuse profiles of H3K27me3 or CTCF. G. Genome-wide fragment cluster score computed at either 1kb shift distance and simulated H3K27me3 at varying shift distances. Our choice for measuring “confinement” can quantitatively distinguish confined versus diffuse experimental ChIP-seq profiles. H. Metaplots showing aggregate depth-normalized H3K27me3 signals from simulated datasets with varying degrees of confinement, with hypothetically no difference in true modification levels at the very center. This reinforces that depth-normalization (e.g., CPM) of a more diffuse profile will yield the impression of a lower peak as compared to confined profile, despite no difference in the absolute value at the center (i.e., a by-product of ChIP-seq depth-normalization). This phenomenon can be important to consider when assessing normalized metaplots I. Confinement scores of H3K27me3 (fragment cluster score at 10kb, see methods) for published ChIP-seq data from the developing mouse brain, ranging from embryonic day 10.5 (E10.5) to birth (P0), in Gorkin et al. (2020)^45^. Diminishing scores indicate the spread of H3K27me3 accompanies early brain development.

**Supplemental Figure SF2.**
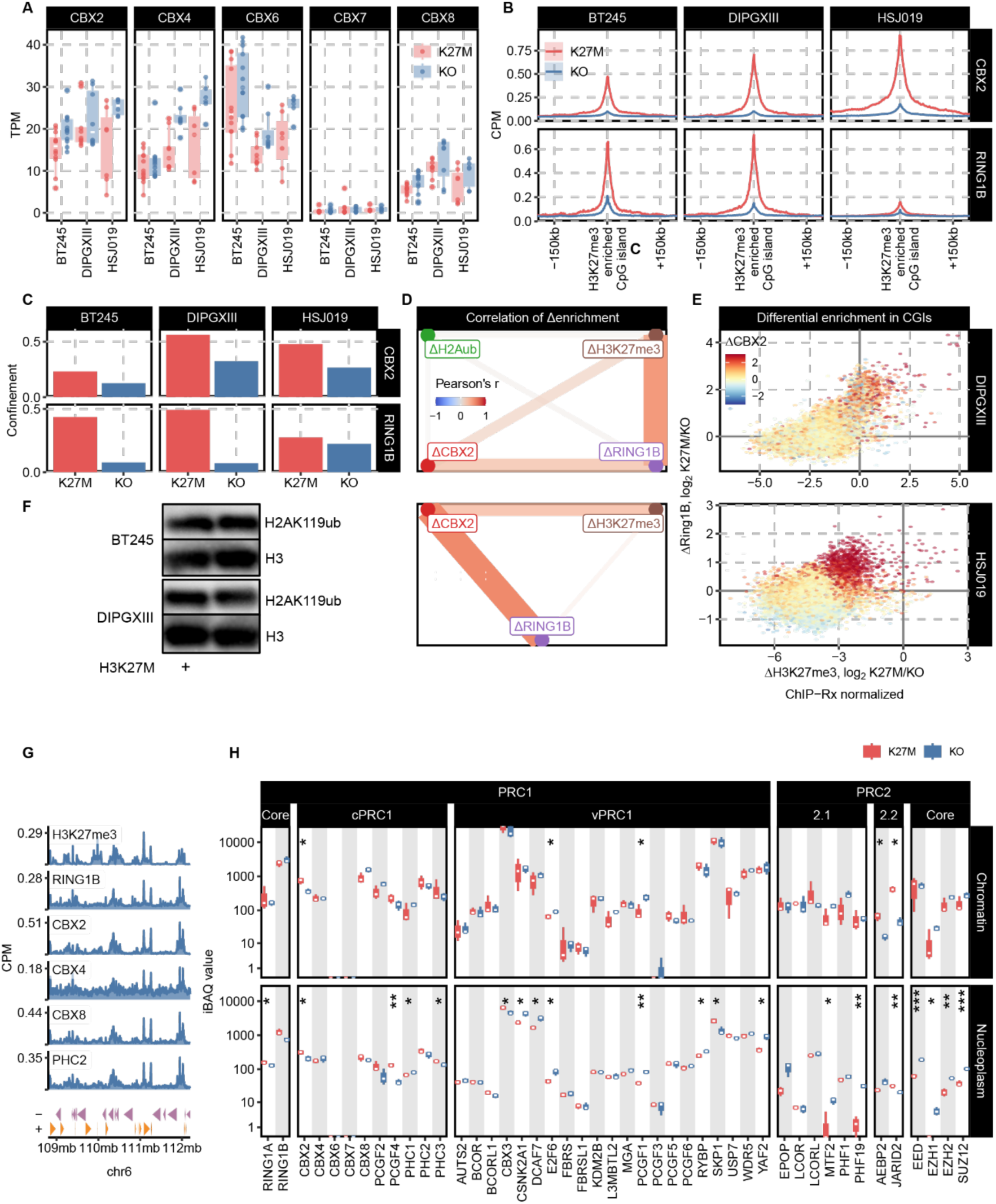
A. Expression of cPRC1 subunit genes (*CBX2*, *CBX4*, *CBX6*, *CBX7*, *CBX8*) in pHGG H3K27M cell lines based on bulk RNA-seq. B. Metaplot of CBX2 and RING1B aggregate ChIP-seq signals around H3K27me3-enriched CpG islands (union set of top 1000 most enriched in both conditions per cell line, as defined previously), normalized by read depth. CBX2 and RING1B occupancy at H3K27me3 sites are consistently diluted by KO of H3K27M. C. Bar graphs showing RING1B/CBX2 ChIP-seq signal confinement scores (fragment cluster score at 10kb, see Methods) in 3 distinct cell lines (BT245, DIPGXIII, HSJ019). RING1B/CBX2 are less confined (ie. more diluted) upon KO of H3K27M mutations. D. Correlation network of differential H3K27me3, RING1B, CBX2 and H2AK119ub enrichment at CGIs of BT245 cells, demonstrating the weak correlation between H2AK119ub changes and the changes of H3K27me3, RING1B and CBX2. Edgewidths reflect the absolute value Pearson correlation coefficients. E. Density plots showing differential CGI enrichment of H3K27me3 (x-axis), RING1B (y-axis), and CBX2 (color code) between H3K27M and H3K27M-KO DIPGXIII (top) and HSJ019 (bottom) cells. Each dot represents a CGI and the differential enrichment is plotted as log2 ratio of K27M/KO. Retainment of H3K27me3 enrichment at CGIs associates with several fold greater enrichment for RING1B and CBX2 ChIP-seq signals, indicating the correlation between H3K27me3 confinement and enhanced cPRC1 recruitment. F. Western blot showing equivalent levels of H2AK119ub abundance in isogenic H3K27M and KO BT245 and DIPGXIII cell lines. G. ChIP-seq/CUT&RUN-seq tracks for H3K27me3 and all cPRC1 subunits profiled, showing that broad domain spreading of H3K27me3 correlates with enrichment of RING1B, CBX2, CBX8 and PHC2 subunits (less so for CBX4) at Mb scale. This indicates cPRC1 can be distributed as both focal peaks and broad domains as determined by the degree of H3K27me3 spreading. H. Mass spectrometry-based measurement of protein abundance (iBAQ) for all subunits of PRC1 and PRC2 complexes, showing most subunits are comparably present in both nucleoplasm (soluble) and chromatin-bound protein fractions of H3K27M and KO cells for the pHGG line DIPGXIII. H3K27M mutations do not therefore dramatically alter the composition or abundance of PRC1/2.

**Supplemental Figure SF3.**
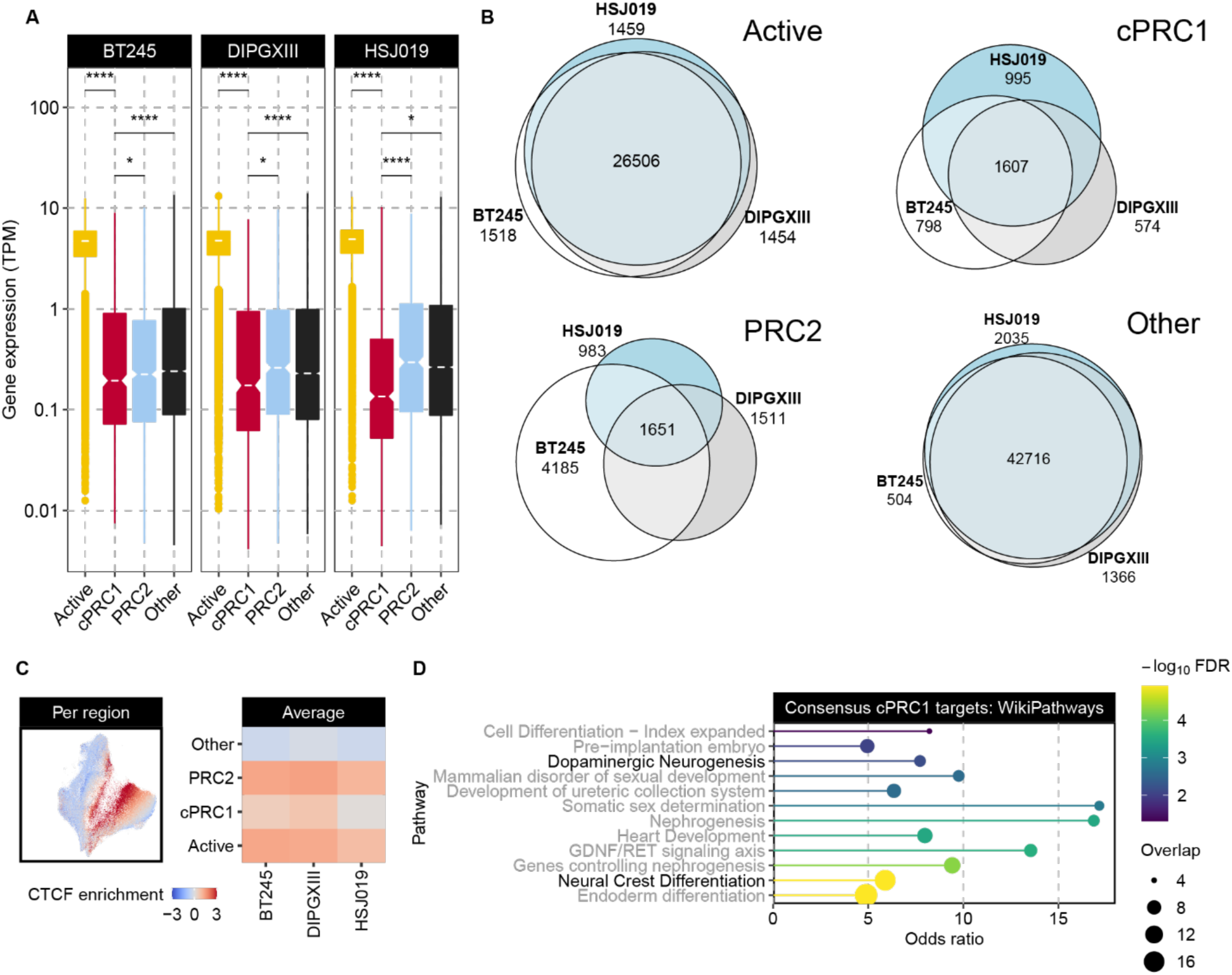
A. Expression of genes (transcripts per million, TPM) associated with the promoters from the four clusters derived in Figure 3, demonstrating the lowest expression levels in the cPRC1 cluster in H3K27M-mutant cell lines. Boxplots’ hinges correspond to the 25^th^ and 75^th^ percentiles, with whiskers extending to the most extreme value within 1.5 × interquartile range from the hinges, whereas the central band mark the median value. B. Euler diagram of sites identified for the four clusters showing concordance of “Active” and “Other” cluster sites among the three H3K27M pHGG cell lines. A substantial fraction of cPRC1 cluster sites also overlap, termed the consensus cPRC1 sites, whereas the PRC2-only cluster sites show less concordance. C. Enrichment of CTCF ChIP-seq signal among the UMAP projection and cluster classification. CTCF is not strongly enriched in the cPRC1 cluster, compared to Active and PRC2-only clusters. D. Enrichr pathway enrichment analysis of consensus cPRC1 targets among three H3K27M pHGG cell lines, demonstrating the enrichment in genes annotated as relating to development and neuron differentiation.

**Supplemental Figure SF4.**
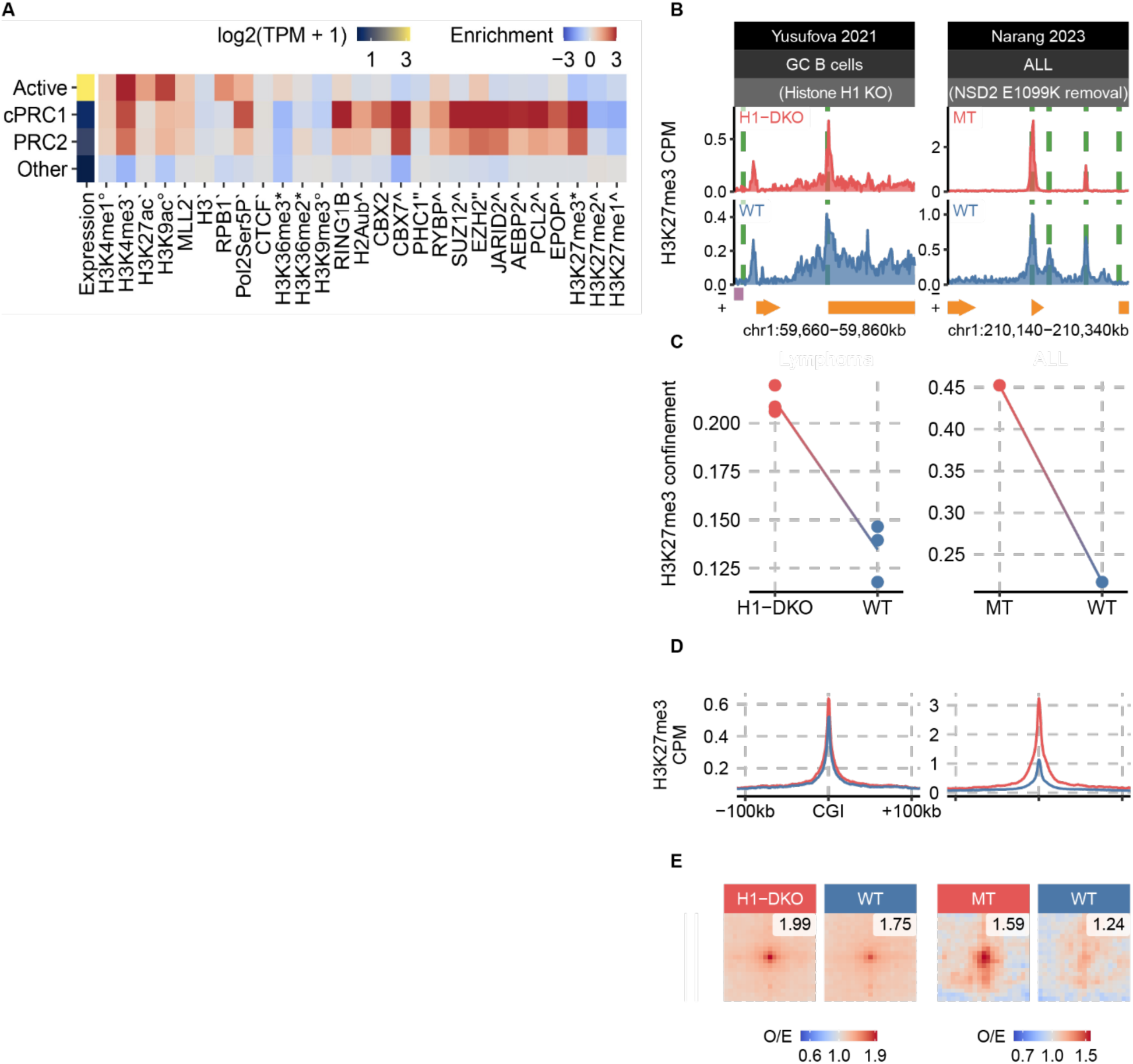
A. Average signals of transcription and chromatin features for CGIs & promoters in each of the four clusters in mESCs (see Figure 4D), demonstrating the characteristic chromatin state of each cluster. Symbols indicate data sources: * = Chen 2022^26^, \ = Kundu 2017^20^, ^ = Healy 2019^71^, ‵ = Mas 2018^72^, ° = ENCODE, no symbol = this study. B. Genomic distribution of H3K27me3 (ChIP-seq coverage tracks in units of counts-per-million-alignments) at representative loci in germinal center B cells and acute lymphoblastic leukemia cells, demonstrating distinctive profiles of confined versus diffuse H3K27me3. C. Measure of H3K27me3 ChIP-seq signal confinement (fragment cluster score at 1kb separation, computed using the tool “ssp”, see methods), comparing confined (H1 KO, NSD2 mutant) versus diffuse profiles. Individual data points correspond to a replicate, with connected points indicating replicates from the same batch; connections not linking points indicate that multiple replicates were sequenced in a batch, and so the links are drawn between the average value per condition. D. Metaplots of H3K27me3 aggregate ChIP-seq signals around H3K27me3-enriched CpG islands, normalized by total read depth. H3K27me3-enriched is defined as the union set of top 1000 CpG islands with the most H3K27me3 alignments in either condition. E. Pile-up of Hi-C interactions among H3K27me3-enriched CpG islands, as defined above, portraying average pairwise contact strength between such regions (in units of enrichment, i.e., observed / expected). Punctate enrichment signal in the center indicates elevated long-range interaction anchored at H3K27me3-enriched CGIs in cells with confined H3K27me3.

**Supplemental Figure SF5.**
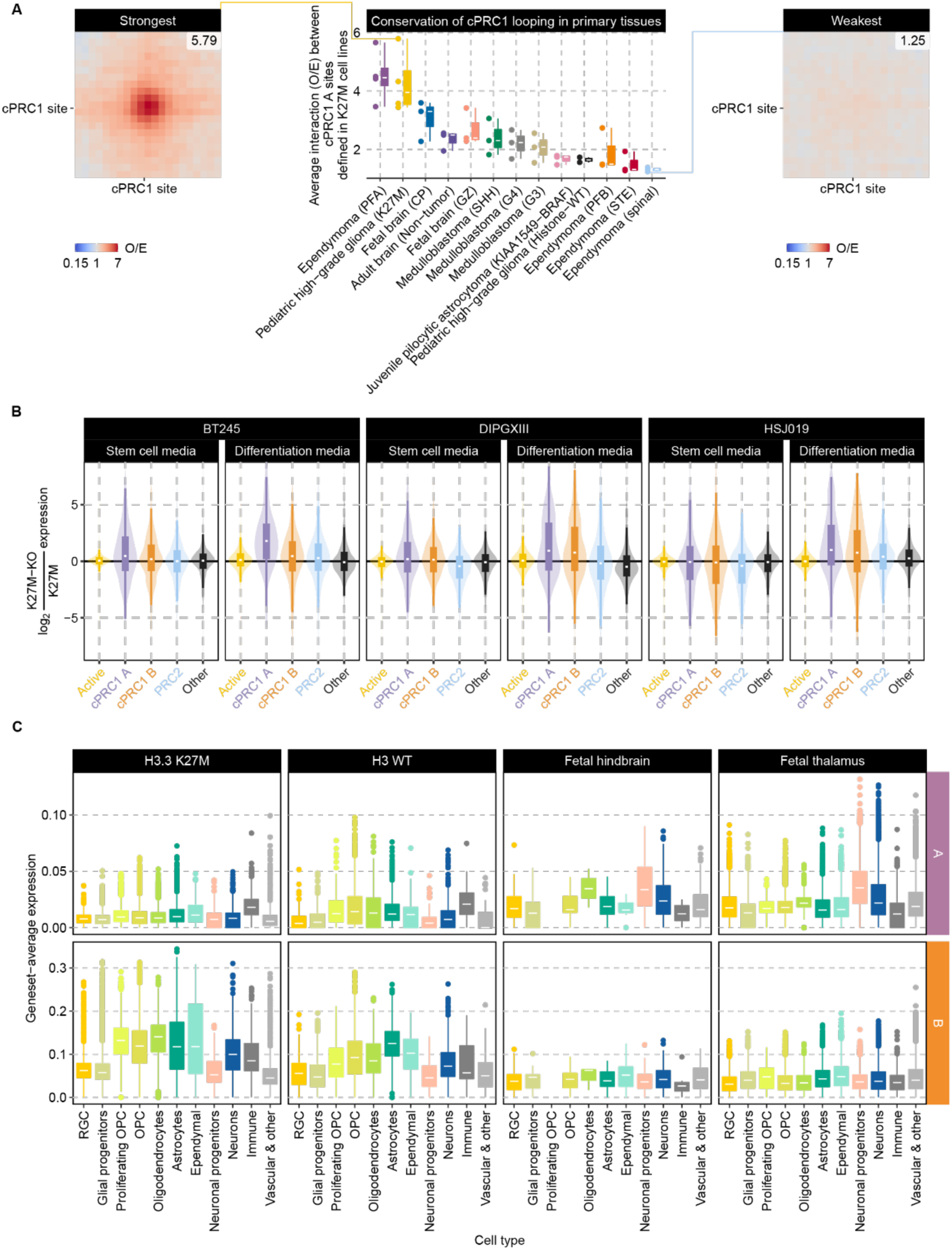
A. Central enrichment (i.e., observed/expected value of the central 3x3 set of pixels) for pile-up of pairwise Hi-C interactions between cPRC1 subcluster A sites in normal or tumor brain tissues samples. The greatest interaction frequencies are found in H3K27M pHGGs and PFA EPNs. Boxplots’ hinges correspond to the 25^th^ and 75^th^ percentiles, with whiskers extending to the most extreme value within 1.5 × interquartile range from the hinges, whereas the central band mark the median value. B. Violin plots showing bulk RNA-seq expression changes, between H3K27M and KO lines in either Stem cell media or Differentiation media, for genes in each cluster. cPRC1 subcluster A genes experience the most consistent transcriptional upregulation upon loss of H3K27M, which becomes more pronounced in Differentiation media condition. Boxplots’ hinges correspond to the 25^th^ and 75^th^ percentiles, with whiskers extending to the most extreme value within 1.5 × interquartile range from the hinges, whereas the central band mark the median value. C. In scRNA-seq datasets, the expression levels cPRC1 subcluster A genes are measured on a cell-by-cell level using single sample Gene Set Enrichment Analysis Score (see Methods for more detail). Distributions of scores across population of cells are plotted, with annotations of cell type population on the x axis (see Figure 5 and Methods for more detail). These scores reveal that cPRC1 subcluster A genes are repressed in a homogenous manner across all cell types in H3K27M pHGG, whereas in WT pHGG and fetal brain tissues, select subpopulations more highly express these genes.

**Supplemental Figure SF6.**
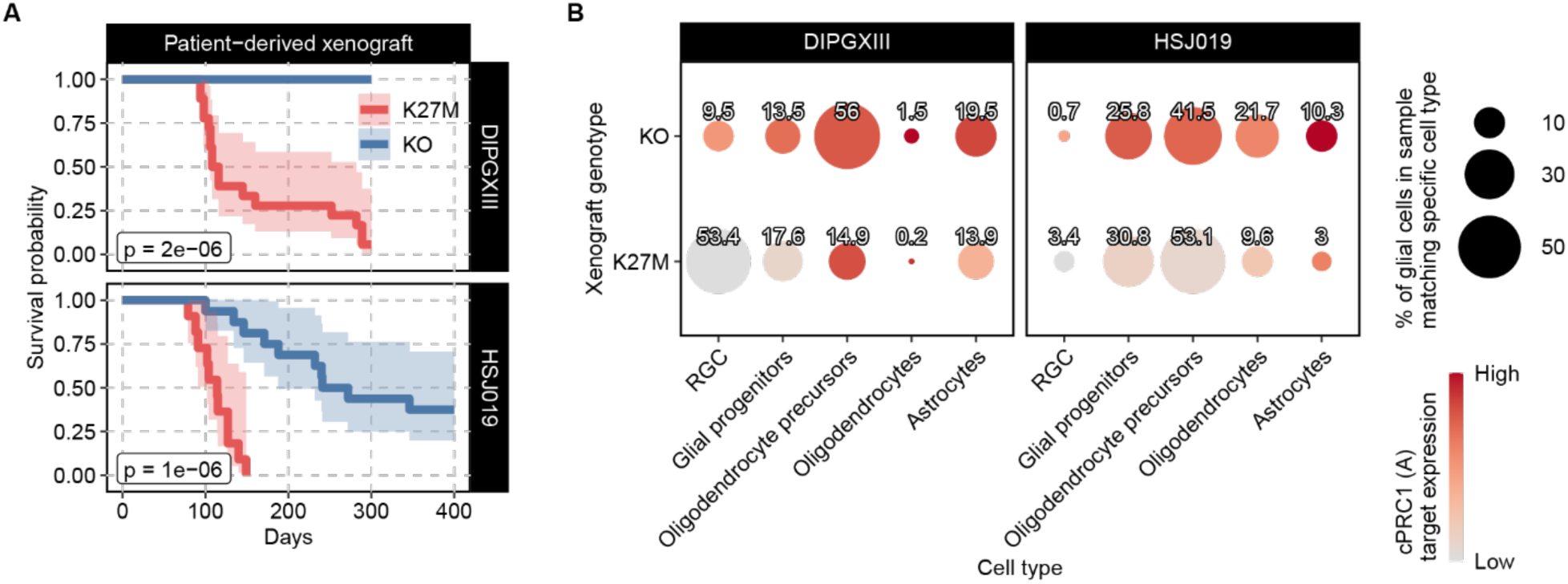
A. Kaplan-Meier survival analysis (with 95% confidence interval in a lighter shade) for xenograft-bearing mice using two other pHGG H3K27M cell lines, displaying loss of tumor formation by H3K27M-KO cells in DIPGXIII, and substantially greater latency and decreased penetrance of tumor formation by H3K27M-KO cells in HSJ019. B. Bubble plots representing fractions of tumor cell populations annotated by cell type classification (bubble size) based on ssGSEA mapping to reference cell types (see Methods). Each bubble is also colored based on the level of cPRC1 subcluster A target gene expression (bubble color). Expression of cPRC1 subcluster A genes is consistently depleted across the spectrum of classified cell types in H3K27M samples, accompanied by lower proportions of more differentiated cell types (astrocytes, oligodendrocytes).

**Supplemental Figure SF7.**
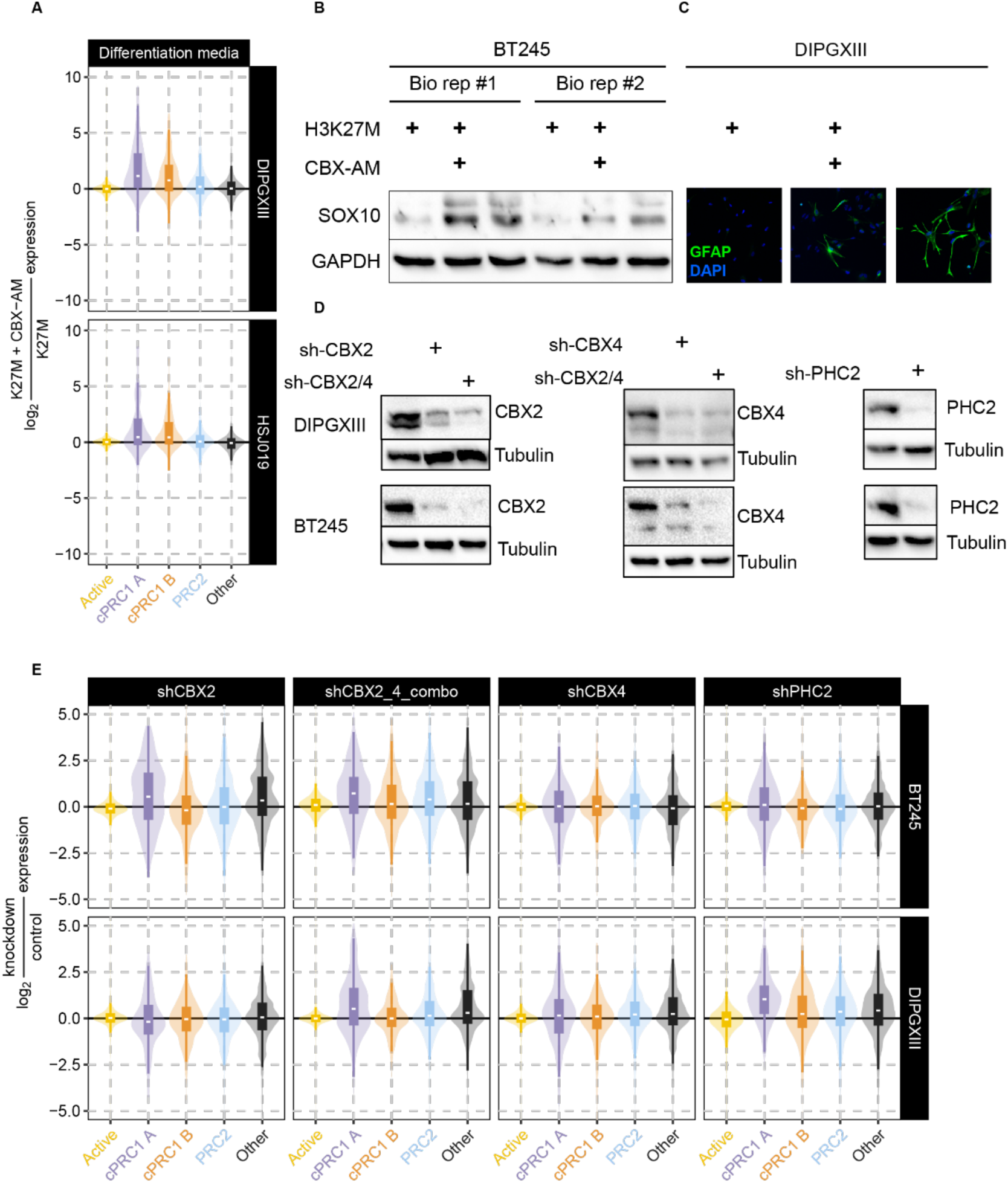
A. Violin plots of genes’ differential expression values in RNA-seq data. CBX-AM treatment results in upregulation of cPRC1 subcluster A genes, and to a less extent subcluster B genes, in H3K27M DIPGXIII and HSJ019 cells treated with differentiation media. Active and PRC2- only cluster genes are not systemically up- or down-regulated due to treatment. Violin plots’ hinges correspond to the 25^th^ and 75^th^ percentiles, with whiskers extending to the most extreme value within 1.5 × interquartile range from the hinges, whereas the central band mark the median value. B. Up-regulation of differentiation marker SOX10 shown by western blot in both H3K27M-KO and CBX-AM treated pHGG line BT245. C. Up-regulation of differentiation marker GFAP shown by immunofluorescence in both H3K27M-KO and CBX-AM treated pHGG line DIPGXIII. D. Western blot validation of depletion of CBX2, CBX4 or PHC2 proteins by lentiviral shRNA delivery to H3K27M cell lines BT245 and DIPGXIII. E. RNA-seq were performed on control and BT245 or DIPGXIII cell lines depleted of CBX2, CBX4, both CBX2 and CBX4, or PHC2 using shRNA knockdown. Violin plots showing bulk RNA-seq expression changes, between control and knockdown lines in Differentiation media, for genes in each cluster. In both cell lines, upregulation of cPRC1 subcluster A genes is observed when both CBX2 and CBX4 are knocked down, indicating combinatorial functions of chromodomain subunits. Violin plots’ hinges correspond to the 25^th^ and 75^th^ percentiles, with whiskers extending to the most extreme value within 1.5 × interquartile range from the hinges, whereas the central band mark the median value.

## Methods

### Patient samples and clinical information

This study was approved by the Institutional Review Board of the respective institutions from which the samples were collected.

### Cell culture

Tumour-derived cell lines were maintained in Neurocult NS-A proliferation media (StemCell Technologies) supplemented with bFGF (10 ng/mL), rhEGF (20 ng/mL), and heparin (0.0002%) on plates coated in poly-L-ornithine (0.01%) and laminin (0.01 mg/mL) (Sigma). Lines were cultured to become differentiated glioma cells by adaptation to media of DMEM-F12 (Wisent) supplemented with 10% FBS (Wisent) for 14 days on coated plates. Measurements of GFAP/SOX10 differentiation in CBX-AM experiments used the following differentiation media; BT245 (DMEM-F12 (Wisent), NeuroCult SM1 Without Vitamin A (StemCell Technologies), N-2 supplement (Thermo Fisher), T3 hormone 3 µg/mL (Sigma), rhPDGF-AA 40 ng/mL (R&D Systems)) or DIPGXIII (DMEM-F12 (Wisent), NeuroCult SM1 Without Vitamin A (StemCell Technologies), N-2 supplement (Thermo Fisher), rhCNTF 50 ng/mL (R&D Systems), rhBMP4 50 ng/mL (R&D Systems)). All lines tested negative for mycoplasma contamination, checked monthly using the MycoAlert Mycoplasma Detection Kit (Lonza). Tumour-derived cell lines were confirmed to match original samples by STR fingerprinting. Parental DIPGXIII and BT245 were respectively obtained from the Monje’s and Ligon’s labs, HSJ19 was derived from a patient sample in the Jabado’s lab. Chromatin extracts from the posterior fossa group A ependymomas (PFA-EP) expressing EZHIP were obtained from the Taylor’s lab. The A1 mouse ES WT and Nsd1-KO lines ^73^ (background C57BL/6 × 129S4/ SvJae F1) were obtained from David Allis lab and maintained on gelatin-coated plates in Knockout DMEM (Gibco) supplemented with 15% ES-cell-qualified FBS (Gemini), 0.1 mM 2-mercapoethanol, 2mM L-glutamine (Life technologies) and LIF. Chromodomain allosteric modulator (CBX-AM) experiments were performed using the compound UNC4976 Trifluoroacetate (Sigma) dissolved in DMSO and applied to cell cultures at 120 µM concentration (RNA-seq differentiation assays for 14 day course of differentiation, Hi-C and ChIP-seq 24 hour treatment). Short hairpin RNA (shRNA) knockdown (KD) cell lines were generated through lentiviral delivery of shRNA pools targeting CBX2 (Origene TL314173), CBX4 (Origene TL314171), or PHC2 (Origene TL310464), followed by puromycin selection at 2 µg/mL for 3 days, comparing matched control lines expressing non targeting shRNAs (sh-NT).

### Mouse orthotopic xenograft

All mice were housed, bred and subjected to listed procedures according to the McGill University Health Center Animal Care Committee and were in compliance with the guidelines of the Canadian Council on Animal Care. Brain tumour cell cultures were transduced with lentiviruses constitutively expressing GFP and luciferase and selected by flow cytometry. Female NSG mice (4-6 weeks) (The Jackson Laboratory, RRID:IMSR_JAX:005557) were used for xenograft experiments using 12-20 mice per experimental group. Cell lines were engrafted using 7×10^5^ cells in the caudate putamen (BT245, HSJ019) or the pons (DIPGXIII) by Robot Stereotaxic machine (Neurostar). Mice were imaged for luciferase signal and monitored for neurological symptoms of brain tumours, including weight loss, epilepsy, altered gait and lethargy. Mice were euthanized when clinical endpoint is reached, followed by removal of the brain. Tissue was sectioned into pieces and a portion of tumour or normal brain was dissociated using the MACS Brain Tumor Dissociation Kit (Miltenyi Biotec). Labelled engrafted GFP+ cells were isolated from dissociated cell population using a BD FACSAria Fusion flow cytometer machine at the McGill University Health Centre Research Institute platform. 5000 viable cells per sample were used as input for scRNA library preparation (see below). Each experimental group was profiled using 3 animals for collection.

### Chromatin immunoprecipitation sequencing

Cells (cell lines or dissociated tumor cells) were fixed with 1% formaldehyde for 10 minutes at room temperature. Cells were lysed in Cell Lysis Buffer for 30 minutes on ice. Nuclei were pelleted at 5000xG for 10 minutes at 4°C, and were resuspended in 200 μL/10 million cells Nuclei Lysis Buffer for 45 minutes on ice. Lysed nuclei were sonicated on a BioRuptor UCD-300 for 18-30 cycles, 30s on 30s off, centrifuged every 15 cycles, chilled by 4°C water cooler. Samples were checked for sonication efficiency to 150-500bp size range by gel electrophoresis. Following centrifugation of samples at 12000xG for 10 minutes at 4°C, supernatants were collected and the chromatin was diluted in RIPA buffer to reduce SDS level to 0.1%. Before ChIP reaction 2% of sonicated drosophila S2 cell chromatin was spiked into the samples for quantification by exogenous reference genome normalization. ChIP reaction for histone modifications was performed on a Diagenode SX-8G IP-Star Compact using Diagenode automated Ideal ChIP-seq Kit. 25ul Dynabeads Protein A beads (Invitrogen) Dynabeads M-280 Sheep anti-Mouse IgG beads (Invitrogen) were washed and then incubated with antibodies (anti-H3K27me3 (1:40, CST 9733), anti-H3K4me3 (1:40, CST 9751), anti-H2AK119ub (1:40, CST 8240), anti-H3K27ac (1:100, Diagenode C15410196), anti-H3K36me2 (1:50, CST 2901), anti-H3K36me3 (1:50, Diagenode C15200183), anti-H3K9me3 (1:50, Active Motif 39161)). 1-2 million cells of sonicated cell lysate combined with protease inhibitors for 10 hr, followed by 20 min wash cycle with provided wash buffers. ChIP reactions for SUZ12, RING1B, CBX2, CBX4, CBX8, SMC1 and CTCF were performed as follows: antibodies (anti-SUZ12 (1:150, CST 3737), anti-RING1B (1:200, Active Motif 39663), anti-CBX2 (Bethyl A302-524 1:200), anti-CBX4 (CST 30559), anti-CBX8 (CST 14696), anti-CTCF (1:400, Diagenode C15410210), anti-SMC1 (1:200, CST 4802)) were conjugated by incubating with 40ul protein A or G beads at 4°C for 6 hours, then chromatin from 5-10 million cells was added in RIPA buffer, incubated at 4°C overnight, washed using buffers RIPA, RIPA+500mM NaCl, LiCl and TE. Reverse cross linking took place on a heat block at 65°C for 4 hr. ChIP samples were then treated with 2ul RNase Cocktail at 65°C for 30 min followed by 2ul Proteinase K at 65°C for 30 min. Samples were then purified with QIAGEN MiniElute PCR purification kit as per manufacturers’ protocol. In parallel, input samples (chromatin from about 50K cells) were reverse crosslinked, and DNA was isolated following the same protocol. Library preparation was carried out using Kapa HTP Illumina library preparation reagents. Half of ChIP DNA was used in End Repair and A-tailing reaction mix, followed by Illumina TruSeq DNA UD Index ligation for 20 minutes at 20°C. The entire bead-purified ligation sample was amplified by 9-12 cycles of PCR. Size selection was performed after PCR using 0.6x/0.8x ratios of AMPure XP beads to collect 250-450bp fragments. ChIP libraries were sequenced using Illumina HiSeq 2000, 2500 or 4000 or NovaSeq 6000 platforms at 50 or 100 bp single reads. For mESC ChIP-seq of RING1B and CBX2, the protocol was adapted from Lee et al. (2006)^74^. For each immunoprecipitation, 30 million cells were dissociated, resuspended in media and crosslinked in 1% paraformaldehyde for 3 minutes at room temperature. Cells were lysed in lysis buffer 1, lysis buffer 2, lysis buffer 3 and 100uL 10% Triton X-100. Samples were sonicated with the Covaris M220 machine (Peak Power 75, Duty Factor 10, 200 cycles for 25 minutes). For antibody conjugation, 75uL of Dynabeads were washed and antibodies (5uL anti-RING1B, Active Motif 39663, or 10uL anti-CBX2 Bethyl A302-524A) were added. Beads were resuspended in lysis buffer 3, and sonicated supernatant was added and rotated overnight at 4°C. Between magnet captures, beads were washed in consecutive buffers; Low salt, High salt, LiCl, and TE with 50mM NaCl. Chromatin was eluted from beads, along with input chromatin sample. Next RNA and protein were digested and DNA was recovered by Qiagen PCR purification kit. Eluted DNA was captured for library preparation and sequencing as previously described.

### Chromatin immunocleavage-sequencing

Reagents and protocol were based on the Epicypher cleavage under targets and release under nuclease (CUT&RUN) commercial protocol. Briefly, 5×10^5^ cells per sample were dissociated, washed and bound to CUTANA Concanavalin A coated Paramagnetic Beads (EpiCypher). Antibodies were bound to cells overnight using 0.5 uL per sample: CBX2 (CST E3N6A), CBX4 (CST 30559), CBX8 (CST 14696), PHC2 (Proteintech 12867-1-AP), RING1B (CST 5694), H3K27me3 (CST 9733), SUZ12 (CST 3737), IgG control (Thermo Fisher 02-6102). Digestion of target chromatin used CUTANA pAG-MNase, followed by DNA collection. Libraries were generated using Kapa HTP Illumina library preparation reagents using 9-12 cycles of PCR, followed by dual 0.6-0.8x size selection using AMPure XP magnetic beads. Libraries were sequenced on the Illumina NovaSeq6000 platform for approximately 30 million reads per library.

### High-throughput chromosome conformation capture

In situ Hi-C libraries were generated from samples, as described previously with minor modifications^42^. Briefly, in situ Hi-C was performed in 7 steps. (1) Cell cultures or frozen tissue were dissociated into single cell suspensions of approximately 2-5 million cells, washed in PBS and crosslinked with 1% formaldehyde for 10 minutes and pelleted. (2) Digestion of DNA used a 4-cutter restriction enzyme (DpnII, 100U) within intact permeabilized nuclei, (3) Filling in and biotinylating the resulting 5′ overhangs and ligating the blunt ends was performed. (4) DNA was sheared to a size of 300-500bp using Covaris LE220 machine (Covaris) at the conditions; Fill Level:10, Duty Cycle: 15, PIP: 500, Cycles/Burst: 200, Time: 58 seconds. Successful shearing was verified using agarose gel separation. (5) Pulling down biotinylated ligation junctions with streptavidin beads was performed. (6) The libraries were amplified from beads in 200 μl of PCR amplification mastermix (KAPA HiFi Hotstart ReadyMix, and KAPA Primer Mix), divided into 50uL aliquots, and run on the program: 98°C 45 seconds, 98°C 15 seconds, 60°C 30 seconds, 65°C 45 seconds, repeating to second step for 8-12 cycles, then 72°C 5 minutes. (7) Amplified libraries were subjected to AMPure XP bead dual size selection from 0.7x-0.9x. Libraries were sequenced for approximately 350M reads using either Illumina HiSeqX PE150 or NovaSeq6000 S4 v1.5 PE150 platforms. Analysis used 2-3 replicates per condition in matched experimental designs, or comparison of brain tumour groups using 3-4 samples per catergory.

### Bulk RNA sequencing

Total RNA was extracted from cell pellets and tumours using the AllPrep DNA/RNA/miRNA Universal Kit (Qiagen) according to instructions from the manufacturer. Library preparation was performed with ribosomal RNA (rRNA) depletion according to instructions from the manufacturer (Epicentre) to achieve greater coverage of mRNA and other long non-coding transcripts. Paired-end sequencing (100 bp) was performed on the Illumina HiSeq 2500 or 4000 platform. Analysis used a minimum of two biological replicates per experimental condition.

### Single cell RNA-seq

The concentration of the single-cell suspension was assessed with a Trypan blue count. Approximately 5000 cells per sample were loaded on the Chromium Single Cell 3′ system (10X Genomics). GEM-RT, DynaBeads cleanup, PCR amplification and SPRIselect beads cleanup were performed using Chromium Single Cell 3′ Gel Bead kit. Indexed single-cell libraries were generated using the Chromium Single Cell 3′ Library kit and the Chromium i7 Multiplex kit. Size, quality, concentration and purity of the complementary DNAs and the corresponding 10X library were evaluated by the Agilent 2100 Bioanalyzer system. The 10X libraries were sequenced in the Illumina 2500 sequencing platform.

### Histone mass spectrometry

The complete workflow for histone extraction, LC/MS, and data analysis was previously described^75,76^. Briefly, cell pellets (∼1 × 10^6^ cells) were lysed, histone precipitated and protein estimated by Bradford assay. Approximately 20 μg of histone extract was then digested with trypsin and a cocktail of isotopically-labeled synthetic histone peptides was spiked in at a final concentration of 250 fmol/μg, followed by propionic anhydride derivatization. nanoLC was performed using a Thermo ScientificTM Easy nLCTM 1000 equipped with a 75 μm × 20 cm in-house packed column using Reprosil-Pur C18-AQ (3 μm; Dr. Maisch GmbH, Germany). Peptides were resolved using a two-step linear gradient from 5 to 33% B over 45 min, then from 33 to 90% B over 10 min at a flow rate of 300 nL/min. The HPLC was coupled online to an Orbitrap Elite mass spectrometer operating in the positive mode using a Nanospray FlexTM Ion Source (Thermo Scientific) at 2.3 kV. Two full MS scans (m/z 300–1100) were acquired in the orbitrap mass analyzer with a resolution of 120,000 (at 200 m/z) every 8 DIA MS/MS events using isolation windows of 50 m/z each (e.g., 300–350, 350–400, …,650–700). MS/MS spectra were acquired in the ion trap operating in normal mode. Fragmentation was performed using collision-induced dissociation (CID) in the ion trap mass analyzer with a normalized collision energy of 35. Raw files were analyzed using EpiProfile^77^. Analysis used a minimum of two biological replicates per experimental condition.

### Western blotting

Histone lysates were extracted using the Histone Extraction kit (Abcam). Whole protein lysates were generated through cell lysis in RIPA buffer and pelleting of non-soluble debris. Lysate protein concentration was determined with the Bradford assay reagent (Bio-Rad). Ten micrograms of protein was separated on SDS-PAGE gels (10% acrylamide) and wet-transferred to a PVDF membrane (GE Healthcare). Membrane blocking was performed with 5% skimmed milk in Tris-buffered saline (50 mM Tris, 150 mM NaCl, 0.1% Tween-20, pH 7.4) (TBST) for 1 hour. Membranes were incubated overnight with primary antibody in 1% skimmed milk in TBST: GAPDH (Advanced ImmunoChemical Inc 2-RGM2 1:1000 dilution), SOX10 (ab212843 1:1000 dilution), CBX2 (CST E3N6A 1:500), CBX4 (CST 30559 1:1000), PHC2 (Proteintech 12867-1-AP 1:1000). Membranes were washed three times in TBST, and the secondary antibody (ECL anti-mouse IgG horseradish peroxidase linked whole antibody) (GE Healthcare) was applied for 1 h in 1% skimmed milk in TBST. Membranes were washed three times and the signal was resolved with Amersham ECL Prime Western Blotting Detection Reagent (GE Healthcare) and imaged on a ChemiDoc MP Imaging System (Bio-Rad). Analysis used a minimum of two biological replicates per experimental condition.

### Immunofluorescence

Cells were plated in a Nunc Lab-Tek II Chamber slide system (ThermoFisher Scientific). Slides were fixed with 4% paraformaldehyde in PBS for 20 min at room temperature, followed by washing three times with PBS. Cells were permeabilized by Triton X-100 (0.05% DIPGXIII, 0.2% BT245), 2% BSA, 5% normal goat serum (NGS) in PBS followed by three PBS washes. Slides were blocked with 5% NGS in PBS for 1 h, followed by overnight incubation with primary antibody in blocking solution: anti-GFAP rabbit monoclonal antibody (CST 12389 at 1:200 dilution), or anti-SOX10 (ThermoFisher Scientific 703439 at 1:400 dilution). Cells were washed three times with PBS and incubated for 1 h with 1:1000 dilution of goat anti-rabbit IgG cross-adsorbed secondary antibody, Alexa Fluor 488 (ThermoFisher Scientific) in blocking solution. Slides were washed three times in PBS and Prolong Gold antifade reagent with DAPI (Invitrogen) was applied. Slides were photographed with a Zeiss LSM780 Laser Scanning Confocal Microscope at ×63 magnification. Each image had the protein of interest (SOX10/GFAP) quantified by fluorescence signal normalized to nucleus count value using ImageJ software. Analysis used minimum of two biological replicates and 5 image replicates per experimental condition.

### Isolation of chromatin-bound protein

This protocol was previously described^78^. Briefly, cells were collected, pelleted, and resuspended in cold E1 lysis buffer (50mM HEPES (Multicell 330-050-EL), 140mM NaCl, 1mM EDTA, 10% glycerol, 0.5% NP-40, 0.25% Triton X-100, 1mM DTT, 1x protease inhibitor tablet (Thermo Scientific (A32955)) using 5 volumes of buffer for 1 volume of cell pellet. The cells were then centrifuged. The supernatant contains the cytoplasm fraction of protein. The pellet was resuspended in the same volume of E1 buffer as previously used, left on ice for 10 minutes and centrifuged. The supernatant was discarded, and the pellet was resuspended in cold E2 buffer (10mM Tris-HCl, 200mM NaCl, 1mM EDTA, 0.5 mM EGTA, 1x protease inhibitor tablet) using 2 volumes of E2 buffer for 1 volume of cell pellet. The cells were centrifuged. The supernatant contains the nucleus fraction of protein. The pellet was then resuspended in the same volume of E2 buffer, left on ice for 10 minutes, and centrifuged. The pellet was then resuspended in cold E3 buffer (500mM Tris-HCl, 500mM NaCl, 1x protease inhibitor tablet) using the same volume as E1 buffer. The cells were sonicated using a Diagenode Bioruptor® Pico Sonication device for 20 cycles (30s on, 30s off) providing the chromatin-bound fraction. All fractions (cytoplasm, nuclear, chromatin) were then centrifuged and the supernatant collected providing the final protein-containing solutions.

### Global quantitative proteomic analysis

For global quantitative proteomic analysis of isolated nucleoplasm and chromatin samples, diaPASEF^79^ based proteomics was used. In brief, samples were denatured in SDC buffer^80^ (1% SDC, 100 mM TrisHCl pH 8.5, and protease inhibitors) and boiled for 15 min at 60°C, 1000 rpm. Protein reduction and alkylation of cysteins, was performed with 10 mM TCEP and 40 mM CAA at 45°C for 10 min followed by sonication in a water bath, cooled down to room temperature. Samples were precipitated with the salt method, as previously described^81^, and pellets were solubilized in SDC buffer (1% SDS /100mM Tris-pH 8.5). Protein digestion was processed for overnight by adding LysC / trypsin mix in a 1:50 ratio (µg of enzyme to µg of protein) at 37° C and 1400 rpm. Peptides were acidified by adding 1% TFA, vortexed, and subjected to StageTip clean-up via SDB-RPS^80^ and dried in a speed-vac. Peptides were resuspended in 10 μl of LC buffer (3% ACN/0.1% FA). Peptide concentrations were determined using NanoDrop and 200 ng of each samples were used for diaPASEF analysis on timsTOFPro. Peptides were separated within 87 min at a flow rate of 400 nl/min on a reversed-phase C18 column with an integrated CaptiveSpray Emitter (25 cm x 75µm, 1.6 µm, IonOpticks). Mobile phases A and B were with 0.1% formic acid in water and 0.1% formic acid in ACN. The fraction of B was linearly increased from 2 to 23% within 70 min, followed by an increase to 35% within 10 min and a further increase to 80% before re-equilibration. The timsTOF Pro was operated in in diaPASEF mode ^79^ and data was acquired at defined 32 × 50 Th isolation windows from m/z 400 to 1,200. To adapt the MS1 cycle time in diaPASEF, set the repetitions to 2 in the 16-scan diaPASEF scheme. The collision energy was ramped linearly as a function of the mobility from 59 eV at 1/K0=1.6 Vs cm^-2^ to 20 eV at 1/K0=0.6 Vs cm^-2^. The acquired diaPASEF raw files were searched with the UniProt Human proteome database in the Spectronaut Pulsar X^82^, a mass spectrometer vendor-independent software from Biognosys. The default settings were used for targeted analysis of DIA data in Spectronaut except the decoy generation was set to mutate. The false discovery rate (FDR) will be estimated with the mProphet approach and set to 1% at the peptide precursor level and 1% at the protein level. The label free quantitation (LFQ) and intensity-based absolute quantification (iBAQ) intensities were further analyzed using the Spectronaut statistical package. Significantly changed protein abundance was determined by unpaired t-test with a threshold for significance of p < 0.05 (permutation-based FDR correction) and 0.58 log2FC.

### LC-MS/MS data analysis

Acquired PASEF raw files were analyzed using the MaxQuant environment V.2.2.0.0 and Andromeda for database searches at default settings with a few modifications^83^. The default is used for first search tolerance and main search tolerance (20 ppm and 4.5 ppm, respectively). MaxQuant was set up to search with the reference human proteome database downloaded from UniProt. MaxQuant performed the search trypsin digestion with up to 2 missed cleavages. Peptide, site and protein false discovery rates (FDR) were all set to 1% with a minimum of 2 peptide needed for identification; label free quantitation (LFQ) and intensity-based absolute quantification (iBAQ) intensities was performed with a minimum ratio count of 2. Match between runs was allowed among each group. The following modifications were used for protein identification and quantification: Carbamidomethylation of cysteine residues (+57.021 Da) was set as static modifications, while the oxidation of methionine residues (+15.995 Da), and deamidation (+0.984) on asparagine were set as a variable modification. Results obtained from MaxQuant, protein groups table was further used for data analysis. LFQ and iBAQ intensities were processed and statistically compared using the Perseus software (version 1.6.15.0).

### ChIP-seq data processing

Raw sequences were first trimmed using fastp v0.22.0 with default settings before alignment using bwa-mem2 v2.2.1 to a combined reference of hg38+dm6 or mm10+dm6 with default settings^84,85^. After identification of duplicates using picard v2.26.2’s “MarkDuplicates” module with default settings, alignments with MAPQ>3 mapping to each species was extracted into separate BAM files using samtools v1.14’s “view” module^86^. Alignments overlapping various kinds of genomic intervals (e.g., uniform 10kb windows, promoters, CpG islands) were subsequently tabulated using bedtools v2.30.0’s “intersect” module^87^. Depth-normalized coverage tracks were generated using deepTools v3.5.1’s bamCoverage module with parameters “--normalizeUsing CPM --centerReads -e 200”^88^. Visualization of aggregate ChIP-seq signal around particular genomic regions was performed using the computeMatrix module of deepTools v3.5.1 in “reference-point” mode with parameters “-bs 1000 -b 1000000 -a 1000000 --referencePoint center”.

### Definition of regions with ChIP-seq enrichment

Regions with focal ChIP-seq enrichment with respect to input controls were identified using MACS v2.2.7.1 with settings “—broad --broad-cutoff 0.1”^89^. To avoid biases, H3K27me3-enriched regions in select panels were alternatively defined as the top 1000 CpG islands with the greatest number of H3K27me3 alignments within 5kb from the midpoint; the union set was then taken between diffuse and confined conditions to further reduce bias towards one condition over another. Comparative ChIP-seq enrichment (e.g., differential CBX2 binding) across conditions were assessed using DiffBind v3.4.0^90^.

### Quantification of ChIP-seq signal confinement

Parameters capturing experimental ChIP-seq data was extracted using the “learn” module of ChIPs v2.4 with default parameters and H3K27me3 ChIP-seq data as well as peak calls from H3K27M pHGG cell line BT245^91^. To simulate spreading, peaks intervals were enlarged from summits by specified widths and then used together with previously learned parameters to generate synthetic ChIP-seq datasets using the “simreads” module of ChIPs v2.4. Signal breadth (“confinement”) of experimental and simulated ChIP-seq datasets was quantified using the fragment cluster score metric from ssp v1.2.1 with default parameters^44^, which captures the degree of clustering for forward-reverse read pairs with specific genomic separation; for the purposes of quantifying H3K27me3 spread, we opted to focus on a separation distance of 10kb based on the typical width of H3K27me3 enriched regions.

### Chromatin-state classification of genomic regions

The union set of GENCODE (v36/vM25) annotated TSSs and mid points of CpG islands from the UCSC Genome Browser were expanded to +/-2.5kb and used as the reference set of regulatory regions for chromatin state classification. The log fold enrichment of ChIP-seq over input for alignments overlapping these intervals were then computed for various targets, producing a table with the rows being promoter / CGI intervals and columns being different ChIP-seq signal sources; this mirrors the expression count matrix of RNA-seq datasets with rows being genes and columns being bulk single cell barcodes or bulk sample identifiers. Analogous to scRNA-seq, dimension reduction was then performed on this data matrix using UMAP with correlation as the distance metric, minimum distance set to 0.01, and neighborhood size varied among (15, 30, 50, 100) depending on the number of datapoints^92^. HDBSCAN was subsequently used for clustering of similar datapoints in UMAP embeddings (of dimensions 5-10) with comparable chromatin states, with minimum cluster and sample sizes varying from (500, 1000, 5000) depending on the number of datapoints^93^. Dimension reduction and clustering was performed iteratively to refine subclusters (e.g., partitioning of all data points into either cluster A and B, and cluster A is then divided into subclusters A1 & A2, and so on).

### Hi-C data processing

Raw sequences were processed using Juicer v1.6 with default parameters against hg38 or mm10^94^. Additionally, Juicer .hic files were converted to cooler .mcool files using hic2cool v0.8.3, after which bias vectors were re-computed using the balance module of cooler v0.8.11 with default settings^95^. To additionally avoid confounding effects contributed by cancer samples with abnormal karyotypes, SV-aware bias vectors were calculated using OneD normalization from R library dryhic 0.0.0.9100 with default settings^96^.

### Global, compartment, and domain level analysis

Compartment scores were computed using the call-compartments module of cooltools v0.4.1 on 100kb resolution OneD-normalized contact maps with GC content as the phasing track. Boundary scores were computed at 50kb resolution using RobusTAD v1.0 with default parameters, taking max (left, right boundary score)^97^. To assess the inter-sample similarity of compartment and boundary scores, genome-wide binned signals per sample were arranged into matrix form (i.e., column = genomic bin, row = sample, value = insulation or compartment score) and used as input for UMAP embedding with correlation as the metric. Global similarity between pairs of Hi-C contact matrices were also directly assessed through HiCRep.py 0.2.6 with “--binSize 50000 --h 5 --dBPMax 5000000”, with the resulting correlation coefficients used for distance matrix computation^98^. Silhouette scores were similarly computed from the same data matrix using inter-sample (1 -Pearson’s r) as the distance, with the goal of identifying which tumor subtype(s) display highly consistent compartment/insulation signatures (or lack thereof).

### Assessment of loop strength

Aggregate peak analysis (APA) / pile-up of interaction profiles was performed using coolpup.py v0.9.5 with default settings to assess the average interaction strength between pairs of genomic intervals such as promoter/CGIs or ChIP-seq peaks^99^. Alternatively, the looping strength of individual pairs of regions (with genomic separation in the range of 20kb-2mb, unless otherwise stated) was measured using chromosight v1.6.1’s quantify module at 10kb resolution with default settings, scoring whether pairs of genomic regions contribute more to “loop-like” patterns on Hi-C contact maps^100^. For promoter/CGI centric analyses, the average chromosight-measured loop score between a given promoter/CGI and neighboring ones (with genomic separation in the range of 20kb-2mb, unless otherwise stated) is taken to assess measures such as intra-class looping strength (e.g., the strength of interaction among cPRC1 targets).

### Differential interactions

Loop score differences between two conditions were calculated based on subtracting chromosight-quantified values for one contact map from another. To independently identify transcriptional regulators whose binding sites strongly overlap with differential loop anchors, BART3D v1.0 was ran for Hi-C maps from each of the three isogenic pHGG cell lines comparing H3K27M vs H3K27M-KO^101^; the ranking (by significance) of transcriptional regulators most strongly associated with differential interaction was then integrated using RobustRankAggreg v1.1 to finally identify which transcriptional regulators were the most consistently enriched across multiple cell lines^102^.

### Bulk RNA-seq data processing

Raw sequences were trimmed using fastp with default settings, after expression quantification was performed using salmon v1.4.0 with default settings against GENCODE annotations (v36/vM25). Transcript-level counts were collated to genes using tximport to produce gene-level count matrices^103^.

### Differential gene expression analysis

DESeq2 v1.34.0 was ran with default settings with gene expression count matrices to identify differentially expressed genes^104^. Gene set enrichment analysis was performed using fgsea v1.20.0 with default settings and taking Wald test statistics as the ranking metric^105^. Gene set over-representation analysis was instead carried out using Enrichr v3.0^106^. Concordance of differential gene expression between different contrasts (e.g., H3K27M vs H3K27M-KO as compared to H3K27M vs H3K27M + CBX-AM) was evaluated using RRHO2 v1.0 with -log_10_(p-value) × sign of logFoldChange as the ranking statistic.

### scRNA-seq data processing

Cell Ranger (10X Genomics, v3.1.0) was used with default parameters to demultiplex and align sequencing reads, distinguish cells from background, and obtain gene counts per cell. Alignment was performed using a joint hg19+mm10 genome reference build, coupled with Ensembl transcriptome build GRCh37 v.82 for hg19 and GRCm38 v.84 for mm10. Intronic counts were excluded. Human cells were extracted if cells were either assigned as human by Cell Ranger or the cell contained greater than 75% of total reads mapping to hg19 in order to obtain adequate numbers of cells per sample. Quality control and normalization was performed using the R package Seurat (v3.1.0)^107^. Cells were filtered based on the following quality control metrics: mitochondrial content (indicative of cellular damage), number of genes and number of unique molecular identifiers (UMIs). Filtering thresholds were set on a per-sample basis where cells were excluded if they had greater than 50% of total reads mapping to mitochondrial read counts, had less than 500 total genes or UMIs, or were outside 2 standard deviations from the mean number of genes or UMIs, respectively. Libraries were scaled to 10,000 UMIs per cell and natural log-normalized. Log normalized counts were used for computing correlations of gene expression and assessing expression of specific genes. Samples were combined by cell line, without any additional transformation of the data.

### Identification of nearest normal cell types in xenograft samples

To assign a nearest-normal cell type to individual cells, Spearman correlation of the log-normalized counts with a reference expression matrix was computed in base R with parameter ‘complete.obs’ to compute covariances. The reference expression matrix was a developmental murine forebrain and pons single cell atlas with average expression values per cluster, as described previously^51^. For each cell, the cluster label with the highest correlation was assigned as the nearest normal cell type.

### Quantification of cPRC1 target expression in single cell data

To assess for enrichment of cPRC1 gene signatures in single cell data, ssGSEA was run to assess enrichment of cPRC1 gene signatures in single cell data^108^, single-sample gene set enrichment analysis (ssGSEA) was run using raw counts per cell and gene sets derived from chromatin-state classification of promoters. Gene sets were derived from chromatin state promoter classification from three separate H3K27M pHGG cell lines, and only consistently classified promoters were kept (i.e., assigned to the same class in all three lines). ssGSEA code was adapted from the GVSA package using parameters ‘alpha = 0.75, normalize = FALSE’^109^. For visualization, proportions were calculated as the fraction of cells of total cells. In cases where only glial cells were visualized, proportions were calculated using fractions of total glial cells.

### PRO-seq library construction

PRO-Seq library construction and data analysis was performed by the Nascent Transcriptomics Core at Harvard Medical School, Boston, MA. Aliquots of frozen (−80°C) permeabilized cells were thawed on ice and pipetted gently to fully resuspend. Aliquots were removed and permeabilized cells were counted using a Luna II, Logos Biosystems instrument. For each sample, 1 million permeabilized cells were used for nuclear run-on, with 50,000 permeabilized *Drosophila* S2 cells added to each sample for normalization. Nuclear run-on assays and library preparation were performed essentially as described in Reimer et al.^110^ with modifications noted: 2X nuclear run-on buffer consisted of (10 mM Tris (pH 8), 10 mM MgCl2, 1 mM DTT, 300mM KCl, 20uM/ea biotin-11-NTPs (Perkin Elmer), 0.8U/uL SuperaseIN (Thermo), 1% sarkosyl). Run-on reactions were performed at 37°C. Random hexamer extensions (UMIs) were added to the 3’ end of the 5’ adapter and 5’ end of the 3’ adapter. Adenylated 3’ adapter was prepared using the 5’ DNA adenylation kit (NEB) and ligated using T4 RNA ligase 2, truncated KQ (NEB, per manufacturers instructions with 15% PEG-8000 final) and incubated at 16°C overnight. 180uL of betaine buffer (1.42g of betaine brought to 10mL) was mixed with ligations and incubated 5 min at 65°C and 2 min on ice prior to addition of streptavidin beads. After T4 polynucleotide kinase (NEB) treatment, beads were washed once each with high salt, low salt, and 0.25X T4 RNA ligase buffer (NEB) and resuspended in 5’ adapter mix (10 pmol 5’ adapter, 30 pmol blocking oligo, water). 5’ adapter ligation was per Reimer but with 15% PEG-8000 final. Eluted cDNA was amplified 5-cycles (NEBNext Ultra II Q5 master mix (NEB) with Illumina TruSeq PCR primers RP-1 and RPI-X) following the manufacturer’s suggested cycling protocol for library construction. A portion of preCR was serially diluted and for test amplification to determine optimal amplification of final libraries. Pooled libraries were sequenced using the Illumina NovaSeq platform.

### PRO-seq data analysis

All custom scripts described herein are available on the AdelmanLab GitHub (https://github.com/AdelmanLab/NIH_scripts). Dual, 6nt Unique Molecular Identifiers (UMIs) were extracted from read pairs using UMI-tools [10.1101/gr.209601.116]. Read pairs were trimmed using cutadapt 1.14 to remove adapter sequences (-O 1 --match-read-wildcards -m). The UMI length was trimmed off the end of both reads to prevent read-through into the mate’s UMI, which will happen for shorter fragments. An additional nucleotide was removed from the end of read 1 (R1), using seqtk trimfq (https://github.com/lh3/seqtk), to preserve a single mate orientation during alignment. The paired end reads were then mapped to a combined genome index, including both the spike (dm6) and primary (hg38) genomes, using bowtie2^111^. Properly paired reads were retained. These read pairs were then separated based on the genome (i.e. spike-in vs primary) to which they mapped, and both these spike and primary reads were independently deduplicated, again using UMI-tools. Reads mapping to the reference genome were separated according to whether they were R1 or R2, sorted via samtools 1.3.1 (-n), and subsequently converted to bedGraph format using a custom script (bowtie2stdBedGraph.pl). We note that this script counts each read once at the exact 3’ end of the nascent RNA. Because R1 in PRO-seq reveals the position of the RNA 3’ end, the “+” and “-“ strands were swapped to generate bedGraphs representing 3’ end positions at single nucleotide resolution. Samples displayed highly comparable recovery of spike-in reads, thus samples were normalized based on the DESeq2 size factors (see stats_summary file). Combined bedGraphs were generated by summing counts per nucleotide across replicates for each condition.

### Visualization

ChIP-seq coverage tracks were imported using rtracklayer and subsequently displayed using ggplot2 v3.3.5^112,113^. Gene annotations were similarly imported and shown through gggenes. Balanced Hi-C matrices were further processed using VEHiCLE with default settings before being imported via RcppCNPy v0.2.10 and similarly visualized using ggplot2^114,115^. 3D structures were predicted from balanced contact matrices using CSynth with default settings^116^. Intersections were shown through Euler diagrams with eulerr v6.1.1.

### Statistical consideration

Unless otherwise stated, Wilcoxon rank-sum tests were used to compare the distribution of metrics between two conditions. When appropriate (e.g., matched isogenic cell lines), a paired instead independent test is performed. P-values are represented as: (****) 0.0001; (***) 0.001; (**) 0.01; (*) 0.05; (ns) 1.

### Public datasets accessed

H3K27me3 ChIP-seq and Hi-C datasets were sourced from: Bonev 2017 (GSE96107)^43^, Gorkin 2020 (ENCODE)^45^, McLaughlin 2019 (GSE124342)^117^, Conway 2021 (GSE162739)^118^, Yusufova 2020 (GSE143293)^49^, Won 2016 (GSE77565)^119^. K27M and K27M-KO pHGG ChIP-seq and RNA-seq data were partially sourced from: Harutyunyan 2019 (GenAP)^39^, Harutyunyan 2020 (GSE147783)^41^, Krug 2019 (GSE128745)^40^. PFA CUT&RUN data were sourced from: Michealraj 2020 (GSE146858)^37^. mESC ChIP-seq data was partially sourced from: Kundu 2017 (GSE89949)^20^, Healy 2019 (GSE127121)^71^, Mas 2018 (GSE99530)^72^, Chen 2022 (GSE186506)^26^. Single cell RNA-seq datasets were sourced from: Jessa 2019 (GSE133531)^51^.

